# Dynamic Loading of Human Engineered Heart Tissue Enhances Contractile Function and Drives Desmosome-linked Disease Phenotype

**DOI:** 10.1101/2020.05.25.111690

**Authors:** Jacqueline M. Bliley, Mathilde C.S.C Vermeer, Rebecca M. Duffy, Ivan Batalov, Duco Kramer, Joshua W. Tashman, Daniel J. Shiwarski, Andrew Lee, Alexander S. Teplenin, Linda Volkers, Brian Coffin, Martijn F. Hoes, Anna Kalmykov, Rachelle N. Palchesko, Yan Sun, Jan D.H. Jongbloed, Nils Bomer, Rudolf. A. de Boer, Albert J.H. Suurmeijer, Daniel A. Pijnappels, Maria C. Bolling, Peter van der Meer, Adam W. Feinberg

## Abstract

The role mechanical forces play in shaping the structure and function of the heart is critical to understanding heart formation and the etiology of disease but is challenging to study in patients. Engineered heart tissues (EHTs) incorporating human induced pluripotent stem cell (hiPSC)-derived cardiomyocytes have the potential to provide insight into these adaptive and maladaptive changes in the heart. However, most EHT systems are unable to model both preload (stretch during chamber filling) and afterload (pressure the heart must work against to eject blood). Here, we have developed a new dynamic EHT (dyn-EHT) model that enables us to tune preload and have unconstrained fractional shortening of >10%. To do this, 3D EHTs are integrated with an elastic polydimethylsiloxane (PDMS) strip that provides mechanical pre- and afterload to the tissue in addition to enabling contractile force measurements based on strip bending. Our results demonstrate in wild-type EHTs that dynamic loading is beneficial based on the magnitude of the forces, leading to improved alignment, conduction velocity, and contractility. For disease modeling, we use hiPSC–derived cardiomyocytes from a patient with arrhythmogenic cardiomyopathy (ACM) due to mutations in desmoplakin. We demonstrate that manifestation of this desmosome-linked disease state requires the dyn-EHT conditioning and that it cannot be induced using 2D or standard 3D EHT approaches. Thus, dynamic loading strategy is necessary to provoke a disease phenotype (diastolic lengthening, reduction of desmosome counts, and reduced contractility), which are akin to primary endpoints of clinical disease, such as chamber thinning and reduced cardiac output.

**Single Sentence Summary:** Development of a dynamic mechanical loading platform to improve contractile function of engineered heart tissues and study cardiac disease progression.

## INTRODUCTION

The heart is responsible for pumping blood throughout the body and it must maintain sufficient cardiac output by adapting to differences in mechanical loads, including stretch during chamber filling (preload) and pressure the heart must work against to eject blood (afterload). These physiological loads are critical to normal heart function, for example, during development they contribute to heart muscle maturation resulting in increased cardiomyocyte alignment and hypertrophy (1). However, preload and afterload can also lead to maladaptive changes in heart muscle, as is the case in hypertension, myocardial infarction and cardiomyopathies (2,3). For example, in arrhythmogenic cardiomyopathy (ACM), mechanical loading on heart muscle that leads to adaptive hypertrophy in healthy patients (i.e. exercise), can accelerate ACM disease progression and lead to ventricular dilation and reduced contractility (4,5). This interplay between patient-specific genetic background and mechanical force is likely critical to understanding a range of cardiac disease states.

Studying the effects of mechanical loading on adaptive and maladaptive changes in heart structure and function is challenging in human patients, driving the need to develop new approaches. Animals have been used to model a wide range of human cardiovascular disease states, but there are inherent differences in physiology, gene expression and pharmacokinetics that limit the ability to replicate complex pathology (6). In vitro 2-dimensional (2D) culture of cardiomyocytes on microengineered surfaces have been used to study structure-function relationships and to create region-specific ventricular myocardium (7,8) Muscular thin films (MTFs), consisting of cells cultured on a flexible film, have enabled these 2D platforms to measure contractility (9,10), and combined with patient-specific human induced pluripotent stem cell (hiPSC)-derived cardiomyocytes can model cardiomyopathies such as Barth’s syndrome (11). However, 2D cardiomyocytes are adhered to a surface (12) and thus do not have the same mechanical cell-cell coupling as found in the native heart (13). To address this, researchers have developed more physiologically-relevant 3D engineered heart tissues (EHTs) in the form papillary muscle-like linear bundles (14–18) myocardium-like sheets(19,20), and ventricle-like chambers (21,22). Incorporating human induced pluripotent stem cell (hiPSC) derived cardiomyocytes, fibroblasts and/or endothelial cells, these EHTs more closely resemble the structure of native myocardium. Further, combined with electrical stimulation to drive contraction, these EHTs can also be functionally matured from an embryonic towards an adult-like phenotype. However, in terms of modeling adaptive and maladaptive loading, current EHTs are limited by anchorage to static posts that result in isometric contraction (i.e. minimal fractional shortening) or incorporation into stretching systems that mimic physiologic strains, but not physiologic loads (27,28).

Here we report development of a dynamic engineered heart tissue (dyn-EHT) platform designed to mimic preload and afterload in order to model adaptive and maladaptive changes in heart structure and function. The goal is to better recapitulate the effect of hemodynamic loading on heart muscle in order to understand how forces alter gene expression, cytoskeletal organization, cell-cell coupling, electrophysiology, and force generation in normal and diseased EHTs. Specifically, we sought to achieve (i) a dynamic preload and afterload that can drive adaptive changes in the EHT, (ii) physiologic fractional shortening during contraction of >10% (i.e., not isometrically constrained), and (iii) a straightforward optical readout of EHT contraction to facilitate experimental throughput. Our results reveal that healthy control embryonic stem cell (ESC) and hiPSC-derived dyn-EHTs show adaptive remodeling under a physiologic loading regime, with structural and functional improvements. Further, dyn-EHTs from patient-derived hiPSCs with a desmoplakin mutation (DSPmut) that leads to clinical ACM, show maladaptive remodeling under the same loading range, with distinct structural abnormalities and functional deficits.

## RESULTS

### Development of the dyn-EHT platform for modeling preload and afterload

To create the mechanical loading platform, we fabricated EHTs around a PDMS strip used to both mechanically load the tissue and measure tissue contractile force. Briefly, human ES or hiPSC cells are differentiated into cardiomyocytes and primary adult cardiac fibroblasts were expanded (**Fig. 1Ai**). At day 0 of the experimental timeline, these cardiomyocytes and cardiac fibroblasts are mixed with collagen and Matrigel, and then cast around elastic polydimethylsiloxane (PDMS) strips immobilized within PDMS wells (**Fig. 1Aii**). Fibroblasts within the mixture then compact the extracellular matrix hydrogel into a linear EHT anchored to the PDMS strip. Cell and matrix compositions were optimized for the formation of a compact tissue with minimal collagen content and high contractility (**fig. S1**). All EHTs are maintained in the PDMS wells in a constrained state until day 14 (**Fig. 1Aiii**). From day 14 to 28, EHTs are either continually cultured in the constrained state or are removed from the wells for dynamic culture (dyn-EHTs) (**Fig. 1Aiv**). In the constrained state, the PDMS strip is immobilized within the well during the entire culture period. This is similar to most current EHT models, where the tissue is unable to significantly shorten, and little active preload is provided to the tissue. In contrast, the dynamic state (dyn-EHTs) allows for the tissue to beat in an unconstrained manner against the PDMS strip, which provides preload between cardiac muscle contractions and allows for increased fractional shortening. Finally, at day 28 both EHTs and dyn-EHTs are removed from culture and functionally assessed based on the ability of the tissue to contract and bend the PDMS strip **(Fig. 1Av**).

**Fig. 1.**
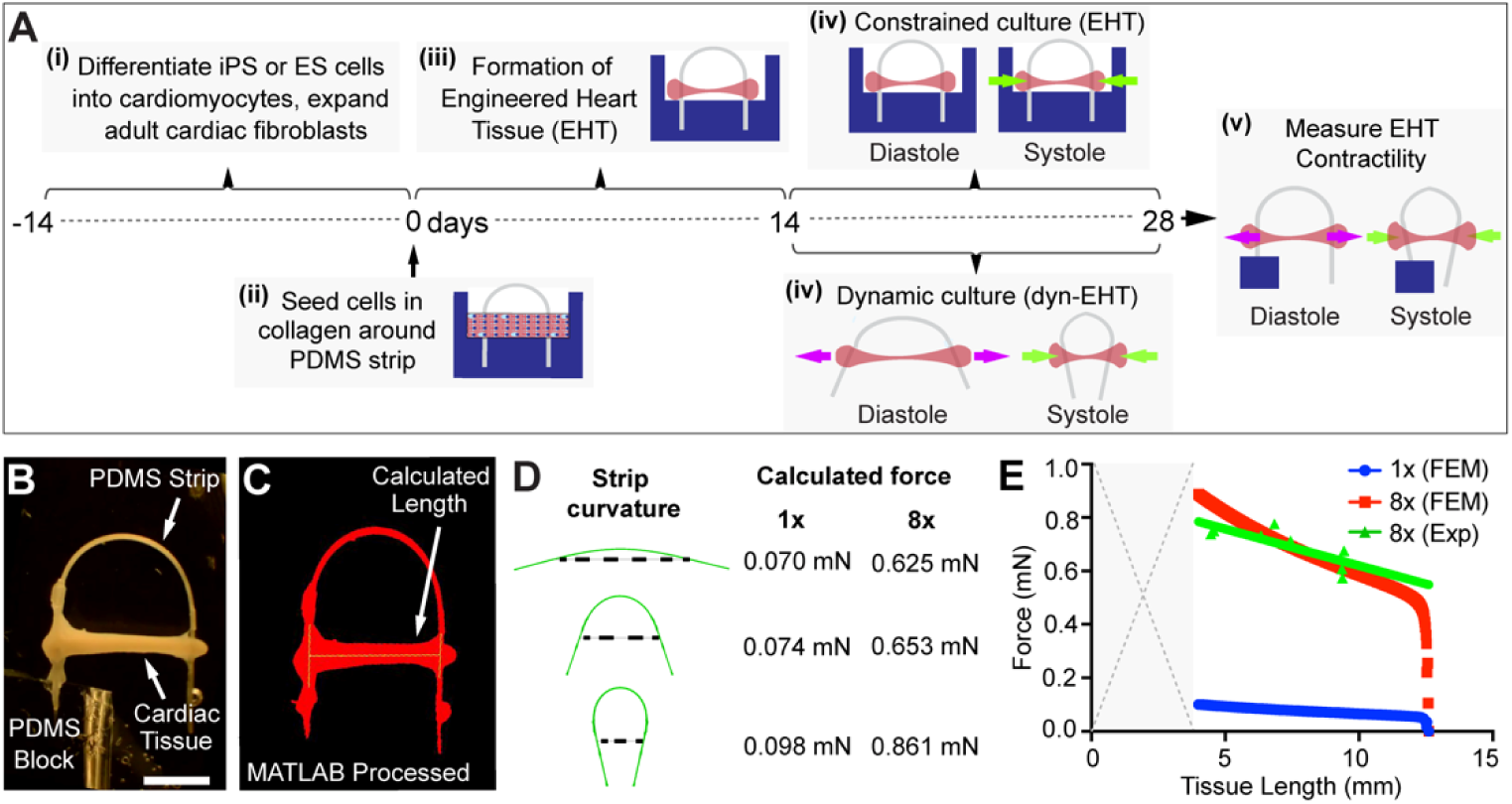
EHTs can be fabricated around PDMS strips for loading and measuring tissue contractile force. (A) Schematic of EHTs casted around PDMS strips. (i) First, cardiomyocytes are differentiated from either ESCs or hiPSCs and cardiac fibroblasts are expanded. (ii) On day 0, cardiomyocytes and cardiac fibroblasts are cast within a Collagen/Matrigel mixture around PDMS strips able to apply two different loads, either 1x or 8x. (iii) From day 4-14, cardiac fibroblasts compact the gel into a linear cardiac tissue between the ends of the PDMS strip. (iv) On day 14 and until day 28, tissues are exposed to constrained (EHTs) or dynamic (dyn-EHTs) loading. Arrows represent the relative amount of preload (pink) and afterload (green) on this tissue within either the constrained or dynamic condition. (v) On day 28, all tissues are measured for contractile force based on the degree of PDMS strip bending. (B) To determine the contractile tissue forces or load that is being applied during diastole, a custom-made MATLAB program is first used to track tissue change in length resulting in PDMS strip bending. (C) Finite element analysis was performed on both 1x and 8x strips to determine the tissue length (represented by the dotted line) needed to induce PDMS strip bending, which could subsequently be used to find the contractile tissue force, or the diastolic load being applied to the tissue. (D) Representative tissue forces with 1x/8x PDMS strip curvature. Dotted line indicates length of the tissue. (E) Average force values obtained at each tissue length with either the 1x or 8x strips. Individual experimental data points are derived from force transducer measurements of strip bending at specific tissue lengths.

Contractility assays are performed in an organ bath where one end of the PDMS strip is mounted in a block and the other end is free to move during each contraction (**Fig. 1B**). A custom MATLAB script is then used to analyze digital video to determine tissue length during contraction, and thus bending of the PDMS strip **(Fig. 1C** and **movie S1**). The force exerted by the tissue is based on known dimensions and elastic modulus of the PDMS strip. We selected strip thicknesses of ∼130 µm (1x) or ∼260 µm (8x), where the thicker strip is an approximately 8-fold increase in load because bending stiffness is proportional to the inverse of the thickness cubed. Finite element modeling (FEM) was used to determine that the applied force could be tuned from 0.05 to 5 mN based on strip dimensions (**fig. S2**), with the 1x and 8x strips being identified as appropriate for these studies having applied forces from 0.07 to 0.9 mN (**Fig. 1D, movie S2**). To validate the FEM, a force transducer was used to measure the force required to bend the PDMS strip. Experimental strip bending forces displayed good agreement with the computational results (**Figure 1E** and **fig. S3**).

### Dynamic 8x loading significantly increases contractile stress generation

At day 28, the contractile function of EHTs cultured under constrained or dynamic conditions and 1x or 8x loading were assessed. There were differences in tissue length and contractility across all four conditions, with a clear improvement in force generation under dynamic 8x loading (**Fig. 2A**). The length of the EHT in the PDMS well during constrained culture was 6 mm, and all EHTs were analyzed out of the well. For the 1x loading, the constrained EHT maintained the same diastolic length as in the PDMS well indicating that the force generated by the 1x strip did not stretch the EHT. Interestingly, for the 1x dyn-EHT the diastolic tissue length actually decreased during culture from days 14 to 28, suggesting that the force generated by fibroblasts compacting the EHT was greater than the force exerted by the 1x strip. The 8x loading results were quite different, with the constrained EHT showing a small increase in diastolic length when removed from the well. However, the dynamic 8x dyn-EHT showed the greatest change, with an ∼50% increase in diastolic length. The tissue fractional shortening showed a similar trend to the diastolic length, with the 8x EHT and dyn-EHT having fractional shortening of 13-15% compared to only 5-8% for 1x loading **(Fig. 2C)**. We found that 1x dyn-EHT had the lowest fractional shortening, suggesting that compaction of the tissue resulted in an increase in tissue stiffness.

**Fig. 2.**
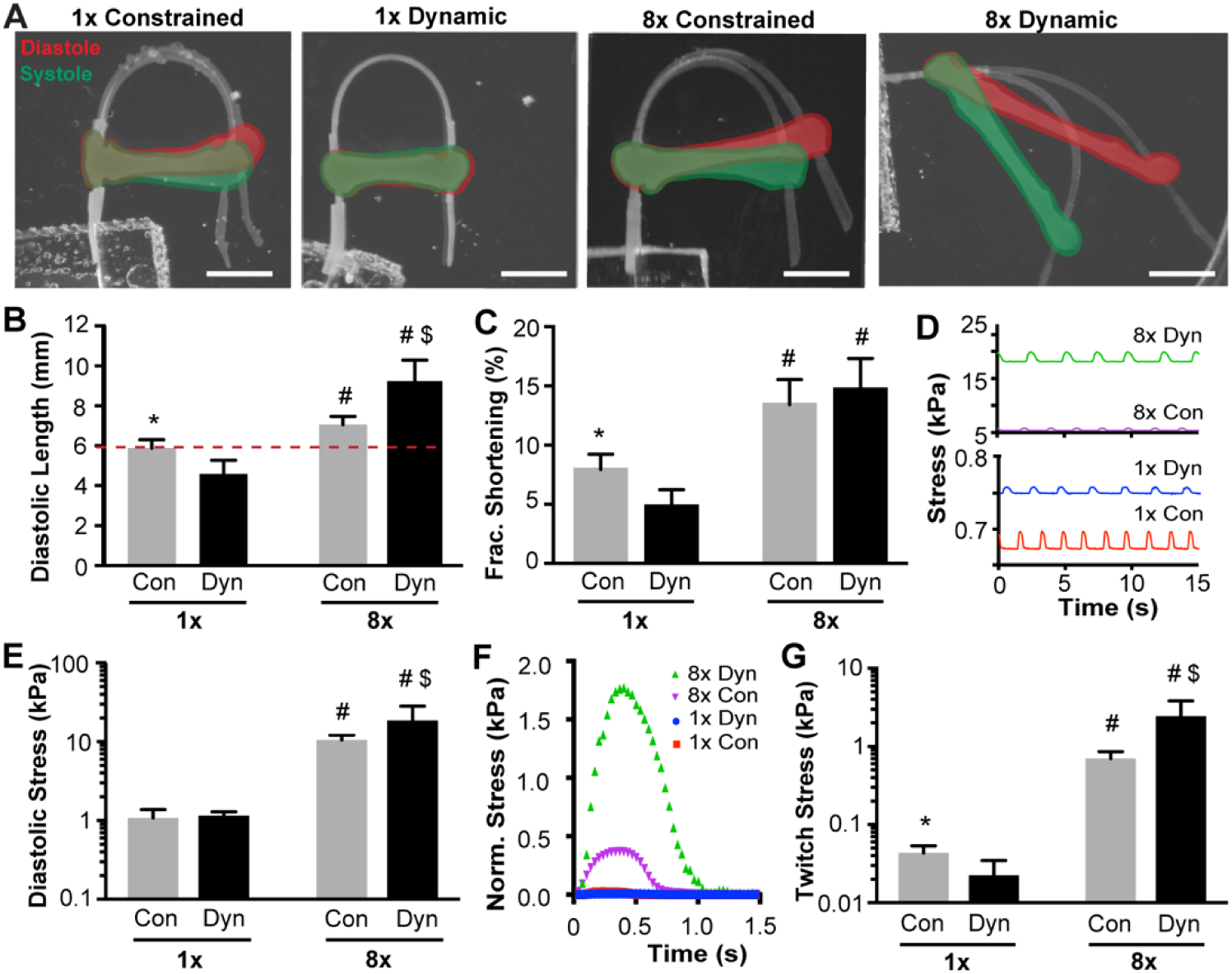
PDMS Strip Can be Used to Mechanically Load Engineered Heart Muscle Tissues. (A-D) Macroscopic tissue contractions overlaid, with tissue in systole (green) and tissue in diastole (red). Scale bars are 2.5 mm. (B) Average diastolic tissue (resting) lengths under each loading condition. The red dotted line indicates the initial length of the tissue while it is immobilized within the PDMS well. (C) Average fractional tissue shortening under each loading type and condition. (D) Cardiomyocyte stress during contraction and relaxation at Day 28. (E) Average tissue diastolic stress at day 28. (F) Representative normalized tissue twitch stresses for one single contraction under each loading type and condition. (G) Average tissue twitch stresses. All graphs show n=5/group for 8x EHT and dyn-EHT, n=6/group for 1x EHT, n=4/group for 1x dyn-EHT. Statistics based on two-way ANOVA with post hoc Holm Sidak test. *p<0.05 vs. 1x dynamic; #p<0.05 vs. 1x dynamic and 1x constrained, $ p<0.05 vs. 8x constrained.

In terms of contractility, the stresses generated by the EHTs were much greater for the 8x versus the 1x loading conditions (**Fig. 2D**). The 8x EHTs displayed elevated levels of diastolic stresses (preload) compared to 1x EHTs (**Fig. 2E**). The elevated diastolic stresses in the 8x group were also associated with higher contractile stresses (afterload). A graph of normalized stress for each condition showing one full contractile cycle demonstrates the relative difference in magnitude in the 1x groups compared to both 8x groups (**Fig. 2F)**. Of note is that the twitch stress (equivalent to the peak systolic stress minus the diastolic stress) observed in the 8x dyn-EHT was more than an order-of-magnitude greater than both 1x conditions and nearly 4-times greater than the 8x constrained EHT (**Fig. 2G**). This result highlights not only the unique impact that dynamic loading has on contractile stress for the 8x dyn-EHT, but also that the magnitude of the load applied to the EHT (1x versus 8x) is important at driving these functional differences.

### Dynamic 8x loading significantly improves maturation state based on myofibrillar alignment, gene expression and electrophysiology

In terms of structure, the EHTs showed major differences in cytoskeletal organization as a function of the different loading conditions. This is important because the myocardium of the vertebrate heart already has highly aligned cardiomyocytes at early time points (29). Similarly, a number of studies using 2D and 3D engineered cardiac tissue have shown that increases in cardiomyocyte alignment are associated with increases in contractile function (7,30–32). Fixing and staining the EHTs for the myofibrils confirmed that all tissues had a dense network of well differentiated cardiomyocytes. However, it also revealed distinct differences in alignment between the 1x and 8x loading conditions (**Fig. 3A**). Quantification of the actin alignment confirmed that there were more myofibrils in the direction of the applied load in the 8x condition, showing the establishment of clear anisotropy. Comparison between the conditions using the orientational order parameter (OOP) further demonstrated that alignment was significantly greater in the 8x versus 1x groups (**Fig. 3B**). While not statistically significant, both qualitative and quantitative analysis of myofibril alignment suggested that the 8x dyn-EHT condition had the greatest degree of anisotropy (**Fig. 3A**,**B**).

**Fig. 3.**
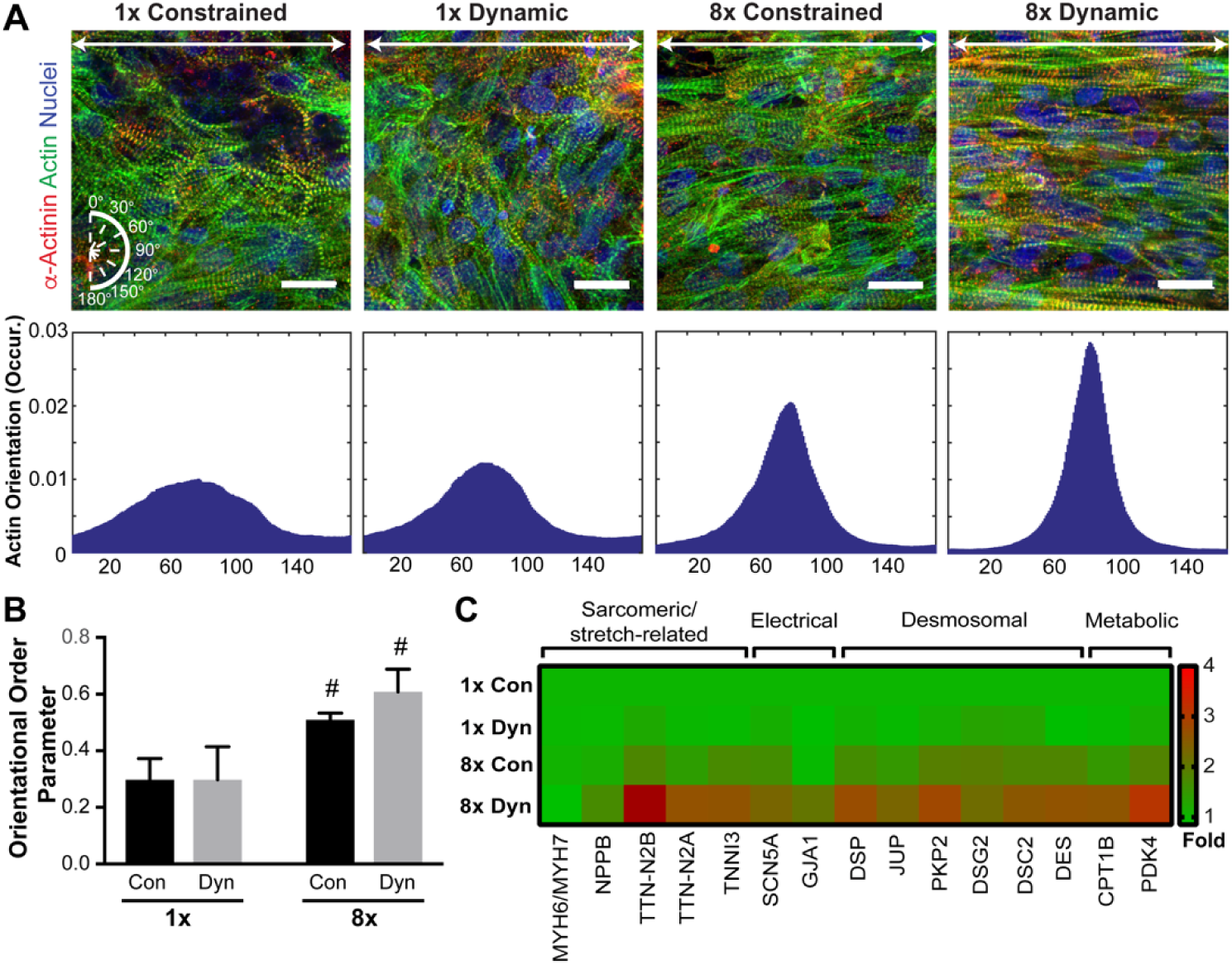
Impact of Loading on Cardiomyocyte Alignment and Cardiac Gene Expression. A) Microscopic images of tissues under each loading condition. The white arrow represents the tissue’s longitudinal axis. Orientation angles were based on white compass (bottom left) with a 90° orientation considered to be along the longitudinal axis of the tissue. Scale bars are 20 µm. Histograms below each image display normalized actin angle occurrence of representative tissues from each loading condition. (B) Average cardiomyocyte alignment (OOP) within engineered heart muscle tissues exposed to different loading conditions. n=3/group. Statistics based on two-way ANOVA with post hoc Holm Sidak test. #p<0.05 vs. 1x dynamic and 1x constrained. (C) Heat map showing log scale geometrical mean of gene expression levels relative to the 1x constrained loading condition. n=3 tissues/group.

Gene expression in the EHTs was also dependent on loading condition. Several genes typically associated with a more mature or adult-like cardiomyocyte phenotype were upregulated in the 8x dyn-EHT (**Fig. 3C)**. We saw an upregulation of all desmosomal genes, including desmoplakin (*DSP*), plakoglobin (*JUP*), plakophilin-2 (*PKP2*), desmoglein-2 (*DSG2*), desmocollin-2 (*DSC2*), and the intermediate filament gene desmin (*DES*), within the 8x dyn-EHT condition. This is in line with previous results, showing that mechanical loading leads to an upregulation of desmosomal genes(15). Metabolic markers associated with fatty acid oxidation (the predominant form of metabolism in the adult heart), including *CPT1B* and *PDK4*, were also upregulated within the 8x dyn-EHT. Further, adult-specific titin (*TTN*) isoforms N2B and N2A, sodium channel *SCN5A* and gap junction Cx43 (*GJA1*) were also upregulated in the 8x dyn-EHT group. While not every gene examined showed an increase in the 8x dyn-EHT, it is evident that a number of genes associated with different aspects of the contractile apparatus in the cardiomyocytes were upregulated.

The electrophysiology of the 8x dyn-EHT condition also showed important difference at both the cellular and tissue scales. At the tissue level, calcium imaging revealed uniaxial conduction along the length of the EHTs from one end to the other (**Fig. 4A**). Quantification of the longitudinal conduction velocity based on the calcium wave propagation showed a statistically significant increase in the conduction velocity of 8x dyn-EHTs compared the 1x conditions from approximately 5 to 10 cm/s **(Fig. 4B)**. This is consistent with the changes in the alignment and gene expression (Fig. 3), as increases in both cardiomyocyte aspect ratio and Cx43 gap junction expression are known to increase conduction velocity (33–36). At the cellular level, sharp electrode measurements of cardiomyocyte resting membrane potential (RMP) were performed on the 1x and 8x dyn-EHTs. The RMP for adult ventricular cardiomyocytes is reported to be in the range of −70 to −90 mV depending on species and measurement technique (37–39). For hESC and hiPSC derived cardiomyocytes, the RMP is typically closer to zero and more fetal or embryonic-like, though a number of studies have reported the ability to use engineered cardiac tissue platforms to produce a more adult-like RMP (24,37,40). We observed that for the 1x dyn-EHT there were two distinct populations, one with a less negative RMP (around −19 mV) and the other with a more negative RMP (around −62 mV) **(Fig. 4C)**. However, for the 8x dyn-EHTs the population of cardiomyocytes with an RMP around −20 mV were absent with an increase in the proportion of more negative RMP cells with an RMP around −60 mV **(Fig. 4D**). This suggests that the increase in applied load between the 1x and 8x strips is able to drive immature cardiomyocytes towards a more adult-like phenotype in terms of RMP.

**Fig. 4.**
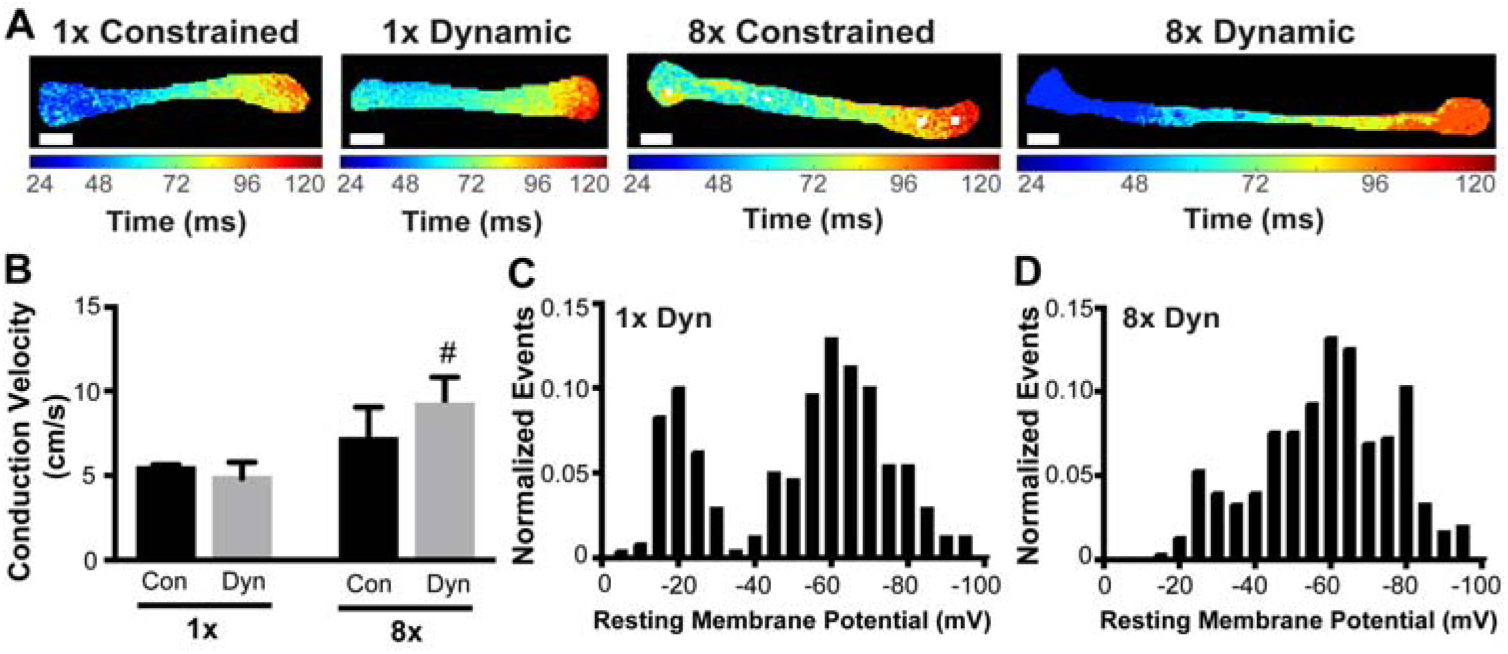
8x Loading Increases Tissue Conduction Velocity and Electrical Excitability. (A-D) Calcium wave propagation across engineered heart muscle tissues under each loading condition. Color map shows time delay of calcium wave as it travels across the tissue. (E) Bar graph of average conduction velocity obtained from calcium transients (F-G) Empirical probability distribution of cells resting membrane potential obtained via sharp electrode measurements from multiple spatial locations (n=3 tissues/group, n=80 cells measured throughout each tissue) within 1x dyn-EHT and 8x dyn-EHT. Conduction velocity based on n=4/ for 8x constrained, 8x dynamic and 1x dynamic groups, and n=3 for 1x constrained group. Statistics based on two-way ANOVA with post hoc Holm Sidak test. # indicates p<0.05 compared to both 1x dynamic and 1x constrained loading conditions.

### Dynamic 8x loading reveals a disease phenotype based on a patient-derived model of arrhythmogenic cardiomyopathy

A key advantage of engineered cardiac tissue is the potential to study human disease states in an *in vitro* system using patient-specific hiPSC-derived cardiomyocytes. As proof-of-concept, we chose to study ACM, where epidemiological data and animal models have indicated that enhanced mechanical loading, i.e. exercise, can accelerate disease progression in patients (5,41–44). Desmoplakin (*DSP*) is an essential component of the desmosome, and pathogenic mutations in *DSP* have been associated with ACM. For this reason, we selected a 50-year-old patient with ACM who was diagnosed with end-stage heart failure and underwent a heart transplantation at the age of 45. The right ventricle of the explanted heart showed fat deposition and fibrosis, and the dilated left ventricle (LV) showed endocardial fibrosis **(Fig. 5A)**. Immunofluorescent (IF) staining of the explanted heart revealed severe reduction of desmoplakin and other desmosomal proteins (**Fig. 5B and fig. S4B**). Next generation sequencing with a targeted gene panel revealed two segregating mutations in *DSP*, where mutation c.273+5G>A was located on one and mutation c.6687delA on the other *DSP* allele (45) (**fig. S4A**). Next, hiPSC cardiomyocytes were generated from this patient (referred to as DSPmut) as well as a control line without the DSP mutation (**fig. S5**). Both of the mutated DSP protein transcripts were functionally evaluated in the hiPSCs. The DSP c.273+5G>A splice site mutation did not result in alternative mRNA splicing, leading to a protein similarly sized as wild type DSP. Nonetheless, protein transcripts from this allele were barely visible on western blot. Mutation c.6687delA, located on the last exon, predicted a frame-shift resulting in a premature stop-codon p.(Arg2229Serfs*32). However, the truncated protein originating from this DSP allele was only detected after inhibition of nonsense mediated mRNA decay p.(Arg2229Serfs*32), indicating that this protein is normally not expressed.(**fig. S6A**). Logically, these hiPSC derived cardiomyocytes displayed reduced desmoplakin IF signal compared to control (**fig. S6B**) and total desmoplakin protein levels were 3-fold reduced in DSPmut cardiomyocytes compared to control (**fig. S6C-D)**. Among other cardiac desmosomal components, levels of desmocollin 2 (DSC2) protein were 2-fold reduced (**fig. S6C**).

**Fig. 5.**
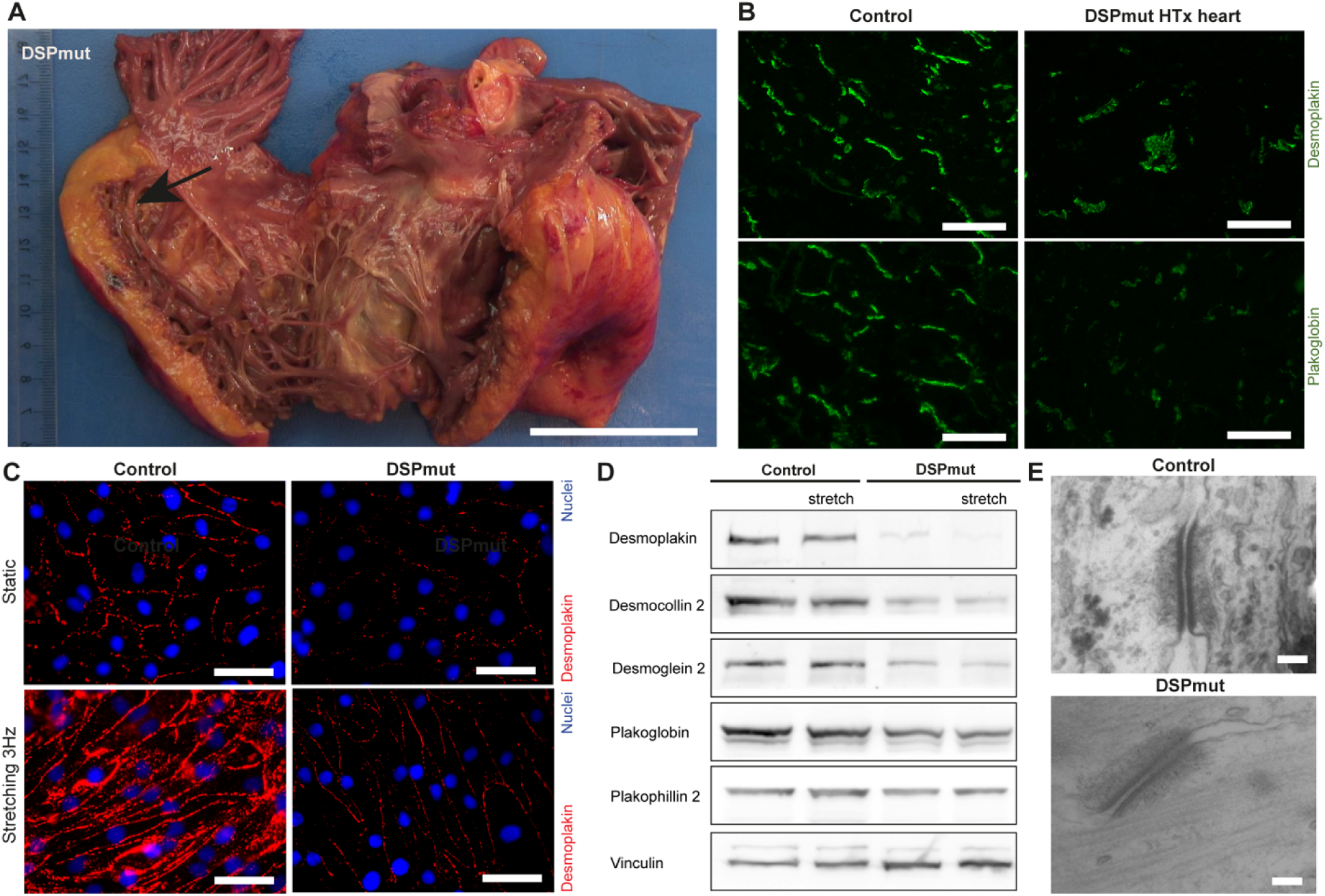
*In Vitro* Phenotyping of DSPmut Cardiomyocytes, as a Case Study of ACM. (A) Explanted heart of the DSPmut patient, where the arrow points to fat deposition. Scale bar is 5 cm. (B) Immunofluorescent (IF) staining of desmoplakin and plakoglobin in transplanted heart (HTx) of DSPmut patient in comparison to a control heart. Scale bars are 50 µm. (C) Desmoplakin localization in DSPmut compared to control cardiomyocytes upon application of 2D stretch. Data is representative for 4 individual experiments. Scale bars are 50 µm. (D) Representative western blot of desmosomal proteins in whole cell lysates after mechanical stretching in 2D. (E) Representative images of desmosomal ultrastructure on TEM. Scale bars are 50 nm.

To understand the impact of 2D versus 3D culture, we first investigated 2D monolayers of hiPSC-derived cardiomyocytes subjected to mechanical stretch. Controls demonstrated a significant increase in localization of desmoplakin, plakoglobin and plakophilin 2 to the cell periphery upon application of stretch **(Fig. 5C, fig. S7, A to B**). However, no increase in desmoplakin localization to the cell periphery was observed in DSPmut cardiomyocytes exposed to 2D stretch (**Fig. 5D**). Despite these significant differences in desmosomal protein localization observed between DSPmut and control cardiomyocytes, whole cell protein levels and mRNA levels were not increased by mechanical stretch applied in a 2D setting (**Fig. 5D** and **fig. S7C**). This is in stark contrast to the upregulation of desmosomal genes observed in 8x dyn-EHTs (**Fig. 3D**). Although a minor increase in the inter-desmosomal space was found, no changes in the number and electron density of desmosomes were observed with 2D stretch (**Fig. 5E** and **fig. S7D**).

Next, we used our 3D system to engineer constrained and dynamic EHTs using 1x and 8x loading. Importantly, there were clear differences for the DSPmut tissues exposed to 8x dyn-EHT compared to the other conditions, with a major increase in tissue diastolic length being observed (**Fig. 6A and Movie S3**). For both 1x and 8x dyn-EHT, IF showed reduced desmoplakin expression in DSPmut EHTs compared to control (**Fig. 6B** and **6C)**. Transmission electron microscopy (TEM) indicated less protein dense desmosomes **(Fig 6D** and **6E and fig. S8A)** and reduced number of desmosomes in DSPmut 8x dyn-EHTs compared to control 8x dyn-EHTs **(fig. S8B)**. These results establish a clear impairment in the desmosomes between the control and DSPmut cardiomyocytes. This is likely the reason why ∼50% of the DSPmut 8x dyn-EHTs broke when removed from the PDMS well at day 14 of culture. The remaining intact DSPmut 8x dyn-EHTs lengthened out significantly more than control 8x dyn-EHTs and any other condition tested (**Fig. 6F**). Fractional shortening was also significantly impacted by loading conditions, with the DSPmut 1x and 8x dyn-EHTs showing a significant decrease from ∼10% to <5% (**Fig. 6G**). Since this was only observed for the dynamic culture condition, but did not depend on load, it suggests that the actual process of undergoing fractional shortening during contraction might somehow degrade the tissue. Finally, DSPmut tissues with 8x loading showed elevated diastolic stresses compared to controls and were similar for constrained and dynamic conditions **(Fig. 6H)**. However, there was a major difference in the twitch stress for the DSPmut tissues between the 8x constrained EHT and the dyn-EHT (**Fig. 6I**). The DSPmut 8x constrained EHT showed a large increase in twitch stress compared to the control EHT suggesting that the increased diastolic stress may initially lead to elevated contractile stresses (afterload). In contrast, the DSPmut 8x dyn-EHT had a significant decrease in contractile force (fig. S9) and twitch stress compared to the control EHT, which makes sense given that the fractional shortening was also impaired. This result demonstrates that the 8x dyn-EHT is uniquely able to manifest a disease-like phenotype from the DSPmut hiPSC-derived cardiomyocytes in the condition with both dynamic culture and higher loading. This strongly suggests that simulating both preload and afterload are required to model certain cardiac disease states such as ACM.

**Figure 6.**
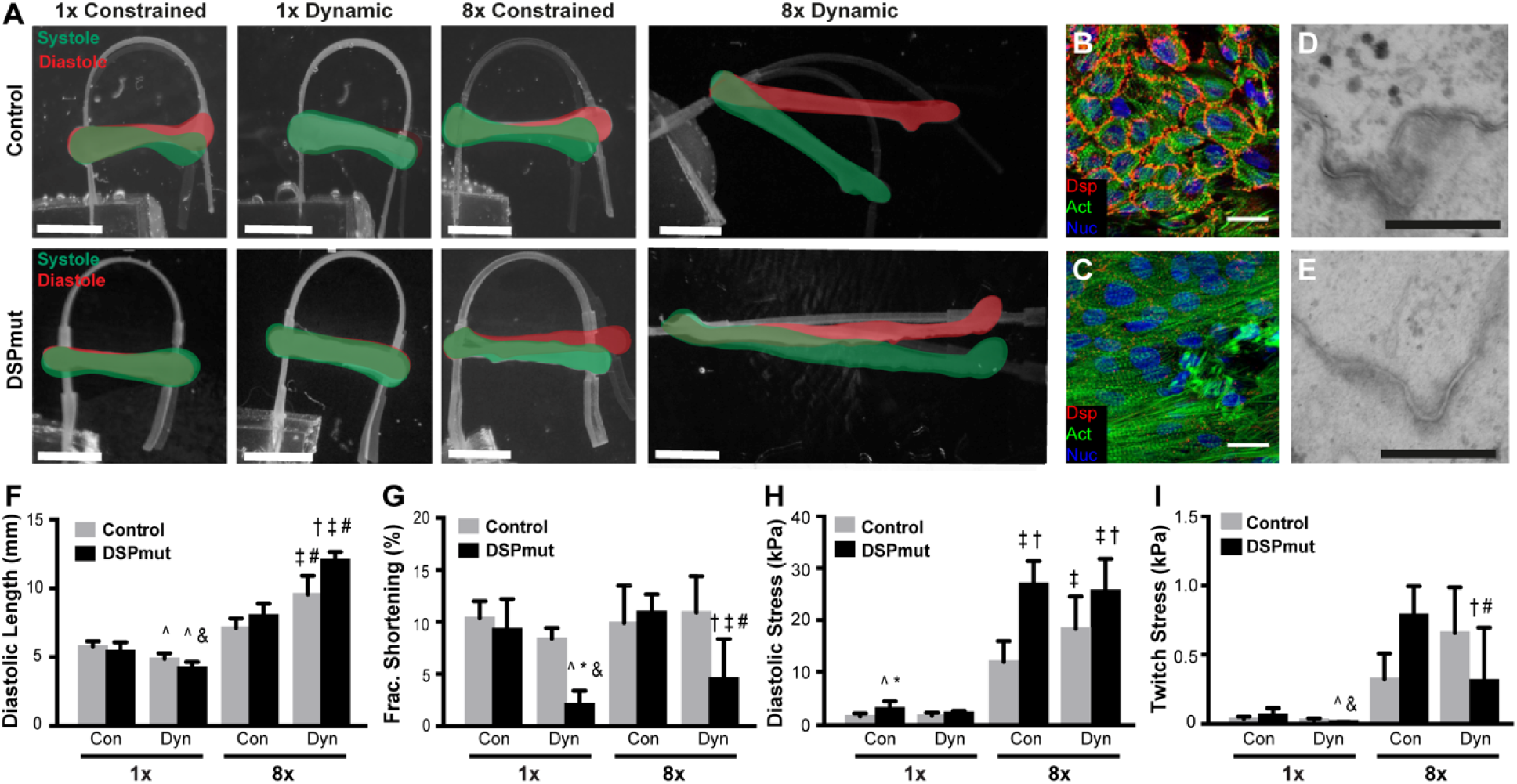
The 8x dyn-EHT model progresses DSPmut Tissues into Disease. (A) Macroscopic tissue contraction of control and *DSP*mut tissues exposed to each loading amount (1x and 8x) and condition (constrained (EHTs) and dynamic (L-EHTs)) taken at day 28. Pictures are an overlay of tissues in systole (green) and diastole (red). Scale bars are 2.5 mm. (B-C) Desmoplakin staining of control (B) and (C) DSPmut 8x dyn-EHTs. Scale bars are 20 µm. (D-E) TEM images of desmosomes of (D) control and (E) DSPmut 8x dyn-EHTs. Scale bars are 300 nm. (F) Bar graph quantifying average diastolic tissue length in control and DSPmut tissues under each loading condition. (G) Fractional shortening at day 28 in control and DSPmut tissues under all loading conditions. (H) Average diastolic stress at day 28 in control and DSPmut tissues under all loading conditions. (I) Bar graph quantifying cardiomyocyte tissue twitch stresses (kPa) at day 28 in control and DSPmut tissues under all loading conditions. Two-way ANOVA with a post hoc Holm Sidak test was performed within each loading condition (either 1x or 8x). ‡ indicates p<0.05 compared to control 8x constrained. † indicates p<0.05 compared to control 8x dynamic. # indicates p<0.05 compared to DSPmut 8x constrained. ^ indicates p<0.05 compared to control 1x constrained. & indicates p<0.05 compared to DSPmut 1x constrained. * indicates p<0.05 control 1x dynamic. % indicates p<0.05 from DSPmut 8x constrained. All graphs show n=8 (DSPmut 1x constrained), n=6 (DSPmut 1x dynamic), n=6 (DSPmut 8x constrained), n=16 (DSPmut 8x dynamic), n=8 (control 1x constrained), n=7 (control 1x dynamic), n=11 (control 8x constrained), n=12 (control 8x dynamic).

## DISCUSSION

The dyn-EHT platform builds off of previous work from our laboratory and other investigators that have used primary, hESC and hiPSC derived cardiomyocytes to engineer functional cardiac tissues in vitro. Indeed, it is critical to acknowledge the important contributions to the field made by researchers using EHTs to model development and disease, and implementing more physiologic-like electromechanical conditioning to drive maturation (14,16-17,20,23,25-26,28,40,61–64). However, these 2D and 3D EHTs are generally isometrically constrained, which means they are unable to undergo significant fractional shortening or be stretched by preload. For instance, a number of 3D EHTs are cast around PDMS or plastic posts or frames that can bend contraction but are otherwise rigid (19,28,46). Constraining the EHT in this way limits the ability to capture changes in tissue length that occur with disease states, such as dilation of the ventricles observed in certain cardiomyopathies(48).

Our results using the dyn-EHT platform shows the importance of modeling aspects of both preload and afterload. We found that dynamic 8x culture of hESC and hiPSC derived dyn-EHTs enhanced contractility, resulting in increased tissue twitch stress, fractional shortening, and cardiomyocyte alignment (Fig. 2). Further, there was an increase in conduction velocity and the proportion of cells with a more negative RMP (Fig. 4) and genes involved in action potential propagation were upregulated, including gap junctions and voltage-gated sodium channels (Fig. 3). It is important to note that though we observed improved function, 8x dyn-EHTs did not reach the functional level of adult human heart muscle. It is possible that by combining our mechanical loading platform with techniques known to improve maturation of engineered heart muscle tissue, like electrical stimulation and exposure to certain biochemical factors such as triiodothyronine (49) and glucocorticoids (50), we may be able to further enhance the function of our 8x dyn-EHTs.While the linear structure and uniaxial stresses in the dyn-EHT are a simplification of the laminar architecture and biaxial stresses in the ventricular wall, we are still able to capture aspects of stress-induced remodeling that have been challenging to observe in other EHT models. In the case of the DSPmut EHT, the dynamic 8x loading resulted in thinning and lengthening of the tissue with increased diastolic stress and decreased twitch stress (Fig. 6). This finding is in line with previous *in vivo* results suggesting that tissue remodeling, including chamber thinning and eccentric dilation, is accelerated with elevated ventricular wall stresses (51,52). In fact, elevated ventricular wall stresses have been proposed to drive the infarct expansion seen post-myocardial infarction (53,54), as well as increase ventricular dilation in dilated cardiomyopathy patients (55–57). Further, strategies aimed at reducing the ventricular wall stress by integrating a temporary patch over the myocardial infarct (58) or injecting materials to thicken the ventricle wall (59) have proved successful to combat the pathological remodeling observed following myocardial infarction. While there have been recent advances in engineering ventricle-like chambers using scaffolds and 3D bioprinting (22,60–62), compared to our dyn-EHT these chambers are much more difficult to fabricate, require a relatively large number of cells to form, and must be connected to a complicated flow system with valves and pressure transducers in order to generate preload.

The dyn-EHT model can also be used to investigate temporal changes in contractility, which is important in order to understand the rate of disease progression. For example, studies have reported an increase in heart muscle contractility when initially exposed to pathological wall stresses (63–65), which is likely due to an overcompensation by remaining heart muscle cells responding to an increase in stretch. Interestingly, we observed a similar finding with DSPmut 8x constrained EHT showing an increase in tissue twitch stress immediately after being removed from the PDMS well (Fig. 6I). Only with prolonged 8x dyn-EHT culture of 14 days did we observe a deterioration in DSPmut tissues, resulting in decreased fractional shortening, contractile force, and tissue twitch stress (Fig. 6F,G,I). While we did not study different time spans of dynamic culture, these pathologic functional changes are likely a direct consequence of the reduced cardiomyocyte coupling (number and density of desmosomes) observed via TEM imaging (Fig 6E). This also parallels heart failure where remodeling of the ventricle occurs prior to decrements in function being observed (51,66).

The ability to mimic disease states using EHTs demonstrates the value of this technology for better understanding the link between genetics and function. Indeed, many studies have already investigated heart diseases based on patient-specific mutations, using either genome-edited or patient-derived hiPSCs. For example, Hinson and colleagues showed sarcomere abnormalities and impaired contractility in EHTs with titin truncating mutations (TTNtv), especially in response to an increased afterload, which was modulated by changing the stiffness of incorporated PDMS posts (46). Wang et al showed reduced contractility in MTFs derived from patients with a genetic defect in Tafazzin, the major phospholipid in the mitochondrial inner membrane (11). Healy and colleagues used variable stiffness microfibers to modulate the afterload on genome edited tissues lacking cardiac myosin binding protein C and found impaired contraction only when tissues were cultured on stiffer fibers (47). The dyn-EHT provides expanded capability by adding preload, which is of particular interest for modeling heart diseases that have a cardiomyocyte junctional or cytoskeletal component, which in fact accounts for a large number of inherited genetic cardiomyopathies (66–69). This is further supported by our data showing that the distribution of mechanical load throughout the cell is different in a dynamic 3D environment when compared either 2D or 3D constrained models (Fig 5 and Fig 6). Importantly, 8x dyn-EHTs formed using control hESC and hiPSC derived cardiomyocytes with no known mutations resulted in upregulation of desmosomal genes, suggesting significant contribution of desmosomes in the adaptive response to mechanical load. In contrast, 8x dyn-EHTs formed using DSPmut cardiomyocytes resulted in tissue lengthening and reduced contractile stress generation. Furthermore, in DSPmut tissues we found reduced amount and expression of desmosomes compared to control, which has also been observed in patients and small animal studies with desmosomal defects(5,71). Taken together, these findings suggest that 3D dynamic mechanical loading of EHTs may be better suited to investigate heart diseases with a cardiomyocytejunctional or cytoskeletal component.

As with any in vitro system, there are a number of important considerations to understand in regard to the dyn-EHT platform. First, in our model the choice of a certain bending stiffness PDMS strip defines the load applied to a tissue. While it is straightforward to change the PDMS strip thickness, length, and width to tune the bending stiffness and applied load (fig. S2), this cannot be changed once integrated into the EHT. A more realistic approach might be to achieve a dynamic change in strip stiffness to change the mechanical load in the same dyn-EHT, similar to what is experienced by the heart during certain disease states. Development of a PDMS strip with a photoactivatable crosslinker that upon illumination would substantially increase the stiffness, is one possible approach. Second, although cardiomyocytes were derived from hESCs or hiPSCs, the cardiac fibroblasts were commercially available and derived from a healthy donor. This was done to minimize variability that can occur between hESC and hiPSC differentiations. Additionally, other cardiac types including endothelial cells and smooth muscle cells could be incorporated in our 3D model but were not explored here and did not appear to be required for the DSPmut condition. Finally, it would be interesting to see if other factors known to influence diseased heart muscle function could be incorporated into our dyn-EHT model. For instance, fibrosis, an important secondary outcome due to injury and remodeling, can reduce compliance of the ventricle and heart muscle fractional shortening (72). Further, it would be interesting to see if the dyn-EHT could be used to model hypertrophic cardiomyopathies, where increased myocardial thickness and lower wall stress would be observed (73). In conclusion, we have developed a model of mechanical loading of EHTs (dyn-EHTs) by incorporating a PDMS strip whose deformation can be used to determine tissue contractile forces but can also be used to mechanically load tissues. We demonstrated that by simply changing dimensions of the PDMS strip, we can modulate the loading on the EHT. Further, because the PDMS strip and EHT can be cultured dynamically, this system allows the tissue to undergo significant fractional shortening and enables the modulation of the preload the cardiac tissue experiences. By incorporating a strip with sufficient load in a dynamic condition (8x dyn-EHTs), we were able to improve the function of control EHTs. Dynamic loading was also important for studying a disease which affects desmosomes in ACM, which is known to be exacerbated by increased hemodynamic loading. Our 3D platform revealed a disease phenotype (tissue lengthening, impaired contractility, fewer desmosomes and reduced electron density), which is similar to the clinical course of disease, including ventricular dilation and reduced cardiac output. Looking forward, our model could have implications for the pharmaceutical industry where patient-derived dyn-EHTs could be used to model heart failure progression in individualized patients with genetic predispositions. This information could be used to test efficacy of drugs that reduce ventricular wall stress, or screen for off-target effects of non-cardiac drugs on heart function. EHTs have also been investigated as a strategy to repair damaged heart muscle *in vivo* (19,68,69) and we demonstrate that both preload and afterload are likely important to improve the function of engineered cardiac tissues prior to implantation.

## MATERIALS AND METHODS

### Study Design

The research objectives prior to this *in vitro* study were 1) to demonstrate that we can modulate the load on engineered heart muscle tissues and 2) to demonstrate improved function over current engineered heart muscle tissue models with minimal preload. We then 3) validated that dynamic mechanical loading is important in creating *in vitro* disease models for studying cardiac diseases. Finally, we compared our disease model to currently used 2D culture techniques. In this study, established stem cells lines, and patient and control derived stem cell lines were generated and subsequently validated. We used multiple lines and multiple differentiations to ensure that a true effect was observed. As our research was performed in two separate research labs, we compared all datasets for proper consistency. Also, for each sub-hypothesis, a minimal of 3 individual experiments were used to calculate statistical significance. Tissues and 2D cultured were excluded if culture conditions differed from our preset timelines.

### Human study approval

This study conforms to the Declaration of Helsinki. Approval of human participants was granted by the medical ethical committee of the University of Groningen (METC 2015.213). Written informed consent was received prior to inclusion of patients. Skin biopsies were gathered for fibroblast culture.

### ESC culture, Cardiomyocyte Differentiation, and Metabolic Purification

Prior to differentiation, ESCs were maintained in Essential 8 (E8) medium (A1517001, ThermoFisher) on Geltrex (A1413202, ThermoFisher)-coated 6 well plates with daily medium changes and passaged at 80% confluence (approximately every 2-4 days). For passaging, cells were washed with 1X PBS and incubated with TrypLE (12604021, ThermoFisher) for 5 minutes at 37C. Cells were then resuspended in DMEM/F12 (11320033, ThermoFisher), centrifuged at 200g for 5 minutes, and replated on Geltrex (A1413202, ThermoFisher)-coated plates in E8 with 2.5 µm ROCK inhibitor, thiazovivin (S1459, Selleck Chemicals). Cardiomyocyte differentiation of human HUES9 ESCs was accomplished by addition of factors that mimic mesoderm induction and cardiac specification during embryogenesis(76–78). On day 0 of differentiation, cells were washed once with 1X PBS and incubated with RPMI/B27 media containing RPMI 1640 (21870076, ThermoFisher) supplemented with 1% v/v L glutamine (25030081, ThermoFisher) and B27 supplement (17504044, ThermoFisher) plus 6 µm CHIR99201 (C-6556, LC laboratories) for 48 hours. On day 2, ESCs were washed with 1X PBS and medium was replaced with RPMI/B27 containing 2 µm Wnt C-59 (S7037, Selleck Chemicals) for another 48 hours. On day 6, medium was replaced with RPMI/B27. On days 8 and 10, medium was changed to CDM3 media, consisting of RPMI 1640 supplemented with 1% v/v L glutamine, 213 μg/mL L-Ascorbic acid 2-phosphate sesquimagnesium salt hydrate >95% (A8960, Sigma), and 500 μg/mL human albumin (A9731, Sigma). On day 12, spontaneously beating cells were passaged for lactate-based metabolic selection of cardiomyocytes(79). To passage for purification, cells were incubated with TrypLE for 15 minutes at 37°C to enable single cell dissociation. Cells were then reseeded in CDM3L, consisting of RPMI 1640 without glucose (11879020, ThermoFisher) supplemented with 213 μg/mL L-Ascorbic acid 2-phosphate sesquimagnesium salt hydrate >95% (A8960, Sigma), and 500 μg/mL human albumin (A9731, Sigma). Cells were maintained in CDM3L for several days before replacing back to CDM3 medium.

### hiPSC Culture, Cardiomyocyte differentiation and Metabolic Purification

Fibroblasts were cultured and reprogrammed to hiPSCs, using nucleofection with three plasmids, pCXLE-hSK, pCXLE-hMLN and pCXLE-hOCT3/4 that were a gift from Shinya Yamanaka (Addgene plasmid # 27078, 27079 and 27076), according to previously reported papers(80,81). Pluripotent colonies were hand-picked and dissociated to single cells using Accutase and TrypLE (Gibco). hiPSCs were seeded, using rock inhibitor (Y27632, Bio-connect) on Geltrex (Thermo)-coated plates, and maintained in E8 medium (Stem Cell). Differentiation was based on slight modifications of previously established protocols (78). Briefly, cells were incubated with RPMI 1640 (ThermoFisher Scientific) supplemented with 5% Knockout™ serum replacement(KOSR, 10828028; Gibco™) containing factors CHIR99201 (13122-1mg, Cayman) and Activin A (120-14E, Peprotech) for 48 hours. On day 2 of differentiation, cells were incubated with medium containing factors Wnt-C59 (5148/10, R&D systems) and BMP4 (SRP3016, Sigma) for 48 hours. Cells were then maintained in previously reported CDM3 medium(82) until day 17. Day 17 post differentiation, cardiomyocytes were treated with dispase and TrypLE to obtain a single cell suspension. Cells were then reseeded for metabolic purification in CDM3L media (lactate-L4263, Sigma(35)) with 2% KOSR. Cells were maintained in CDM3L for several days before replacing back to CDM3 medium. Cardiomyocytes were then used for 2D and 3D experiments. To investigate the effect of *DSP* mutation c.6687delA on protein expression, nonsense mediated decay inhibition was performed on cardiomyocytes using 50 µM NMDi14 (previously mentioned(83), SML1538-5MG, Sigma) for 6 hrs.

### Human Cardiac Ventricular Fibroblast culture

Human Ventricular cardiac fibroblasts (NHCF-Vs) were obtained from Lonza and expanded maximally to ensure use of a single cell source throughout our experiments. Cells were maintained in FGM-3 Medium (CC-4526, Lonza) and passaged at 80% confluence.

### Fabricating Cardiac Tissues Around PDMS Strips

To fabricate engineered heart muscle tissues around strips, first PDMS wells were created. To create the wells, molds were designed in Solidworks and printed using either Verowhite plus RGD835 using an Objet Connex350 (Stratasys) or Dental SG resin (RS-F2-DGOR-01, Formlabs) on a Form 2 SLA printer. Polydimethylsiloxane (PDMS) wells were created to cast the tissue around the strip by casting Sylgard 184 PDMS (Dow Corning) in a 10:1 base to curing agent ratio. Sylgard 184 was mixed prior to casting using a Thinky conditioning mixer (Phoenix Systems) for 2 minutes at 2000 RPM each for mixing and defoaming. Molds were degassed for 30 minutes under vacuum to remove bubbles, cured at 65 °C for 4 hours, and then PDMS wells were removed from the molds. Subsequently, PDMS-strips were laser-cut from PDMS sheets (Bioplexus, Inc), either ∼130 µm (SH-20001-005) or ∼260 µm (CUST-20001-010) thick, using a ProtoLaser U3 (LPK5) laser cutter. Because bending resistance of the strip is proportional to the thickness-cubed, a profilometer was used to determine each strip thickness (VHX-5000).

Prior to cardiac tissue fabrication, wells were sterilized via sonication in 50% v/v ethanol for 30 minutes and PDMS strips were sterilized with UV-Ozone treatment for 15 minutes. Sterile PDMS wells were fixed to one well of a 6 well culture plate using a thin layer of vacuum grease. PDMS wells were then incubated with 1% w/v Pluronic F-127 (P2443, Sigma) for 10 minutes to reduce collagen attachment to the well and washed three times in 1X PBS. Sterile PDMS strips were then placed in the slits in the bottom of the well to secure the strip during cardiac tissue casting. Collagen I derived from rat tail (354249, Corning) and Matrigel (354263, Corning) were used to fabricate 3D cardiac tissues around the PDMS strips. The final concentrations were 1 mg/mL Collagen I, 1.7 mg/mL Matrigel, 10% 10X PBS, 2.3% 1N NaOH, 18.75×10^6^ total cells/mL consisting of 90% cardiomyocytes and 10% cardiac fibroblasts. Ingredients were mixed with a pipette and then 80 µL of the cell mixture was pipetted into each well around the PDMS strip. 6-well plates containing the molds and cell suspension were then placed at 37°C to allow for gelation. Following a 45-minute incubation period, cardiac tissues were incubated and maintained in heart muscle tissue media containing RPMI-1640 supplemented with 1% v/v KnockOut™ Serum Replacement.

### Cardiac Tissue Loading

To mechanically load the tissues, strips of two different thickness’ (either ∼130 µm or ∼260 µm) were incorporated into the cardiac tissues during tissue formation, where the thicker strip represented an approximate 8x increase in load being applied to the tissue. On day 14, tissues were exposed to either constrained or dynamic mechanical loading from day 14 to day 28. In the constrained condition, the strip was held immobilized within the well throughout the entire culture period. In contrast, the dynamic condition allowed for the tissue to beat in an unconstrained manner against the PDMS strip. For dynamic loading, a PDMS block with a vertical cut was adhered to a new well with vacuum grease and one end of the strip was fixed into the vertical cut in the PDMS block. Tissues were maintained in heart muscle tissue media in either constrained or dynamic culture until day 28.

### Calcium Imaging and Conduction Velocity Measurements

To assess tissue conduction velocity, engineered heart muscle tissues were stained with fluo-4 (1 µm), a calcium indicator dye. An excitation-contraction decoupler, blebbistatin (10 µm) was used to prevent tissue contraction, which would make it difficult to perform automated analysis on the calcium wave front. Calcium imaging (50 frames per second) was performed using a Prime 95B Scientific CMOS camera (Photometrics) mounted on an epifluorescent stereomicroscope (Nikon SMZ1000) with a GFP filter and an X-cite Lamp (Excelitas). Conduction velocity (cm/s) was calculated by obtaining the average distance of the tissue in centimeters and dividing that by the average time (in seconds) it takes for the wave to traverse the tissue. For activation maps, calcium transients from spontaneous contractions were then post-processed with a custom-made MATLAB program using calcium signal peak times and cross-correlation.

### Tissue Contractility Assay and Measurements

During engineered heart muscle tissue contraction, the tissue decreases in length, resulting in PDMS strip bending. Based on known dimensions and elastic modulus of the PDMS strip, the degree of strip bending can be used to determine the force the tissue is exerting during contraction. To determine tissue twitch stress, tissue diameter was measured, and cross-sectional area was estimated assuming tissue area conformed to a circular cross-section. Twitch stress measurements were found by dividing average twitch force by average cross-sectional area of cardiomyocytes within the tissue. To image PDMS strip bending due to cardiac tissue contraction, a PDMS block with a vertical cut was adhered to a petri dish using vacuum grease and one end of the strip was fixed into the vertical cut in the PDMS block. During the contractility assay, tissues were maintained in Tyrode’s (T2145, Sigma) solution on a custom-heated stage at 37°C +/-1°C. The PDMS strip was imaged with a Nikon DSLR camera mounted on a Nikon SMZ1500 stereomicroscope to visualize strip bending due to cardiac tissue contraction. A custom-made MATLAB program was created to automatically calculate the change in length of the tissue from contraction videos (fig. S8). Look-up tables relating tissue length to force required to induce PDMS strip bending were created using a finite element modeling. Look-up tables were derived from finite element modeling of strip bending performed in ANSYS software (fig. S9). Modeling was performed by applying known displacements to the PDMS strip and then obtaining the reaction force associated with these displacements. The PDMS strip was modeled as a 3D deformable, extruded solid with two different thicknesses. The strip was considered linearly elastic, isotropic, and incompressible with a Young’s modulus of 1.59 MPa for ∼130 µm strips and 1.89 MPa for ∼260 µm strips, respectively. Elastic moduli were derived from tensile testing of laser cut, dogbones derived from each respective material. Briefly, dogbones mounted on an Instron 5943 and stretched at a rate of 1 millimeter/minute. The average cross-sectional area of the inserts was manually determined prior to testing and dogbone thickness was determined with profilometer (VHX-1000) measurements. Stress was calculated by dividing the force by the cross-sectional area of the strip. Stress was then plotted against the strain and slopes of the individual curves in the linear region of the stress-strain graph (0-10%) were averaged to obtain the elastic modulus of the PDMS.

### Sharp electrode measurements

Tissues were washed 5x times with Tyrode’s solution. Glass microelectrodes with tip resistances of 8-20 MΩ when filled with 3 M KCl were used to impale the tissues. Measurements were performed using a MultiClamp 700B amplifier and analogue signals were low-pass filtered (10 kHz) and digitized at a sample rate of 20 kHz via a Digidata 1440A A/D digitizer (both Axon Instruments/Molecular Devices, Union City, CA). All recordings were performed at room temperature. Data was analyzed using pCLAMP 10.7 acquisition software (Axon Instrument), Excel (version 2010, Microsoft, Redmond, WA) and Prism 7.02 (GraphPad Software, La Jolla, CA). Both datasets were normalized (by dividing the number of events in each bin by the total number of events). Only measurements with stable resting membrane potentials were included in this study.

### Gel electrophoresis and western blotting

Cellular proteins from cells or tissues were extracted using a buffer containing 62 mM Tris-HCl, 2.5% SDS and 1 mM EDTA. Before extraction protease inhibitor (Roche 11873580001), phosphatase inhibitor cocktail 3 (p2850; Sigma) and Sodium orthovanadate were added. After protein quantification (Pierce), sample buffer was added (final solution contains 10% glycerol, 5% β-Mercaptoethanol and bromophenol blue). Samples were heated at 99°C before loading. Proteins were separated using SDS-PAGE gels and transferred to PVDF membranes using semi-dry or tank blotting. Membranes were blocked using 5% ELK and incubated on a shaker overnight at 4°C with primary antibody (Table S1) and 1hr on a shaker at room temperature with secondary HRP-labeled antibodies. As loading control, either GAPDH or vinculin was used depending on the protein’s molecular weight. For detection, electrochemiluminescence was used. Protein quantification was done with the ImageJ gel analyzer.

### Immunofluorescence

For whole mount imaging, cardiac tissues were fixed in 4% formaldehyde overnight and permeabilized with Triton-X 100 for one hour at room temperature. 2D cultured cells were fixed and permeabilized with ice cold methanol:acetone for 10 min. *Ex vivo* tissues were cut with a cryostat and air dried before staining. Tissues/cells were incubated with BSA/PBS blocking buffer containing serum of secondary antibodies’ host. Tissues/cells were then incubated with primary antibodies in blocking buffer accordingly. Following primary antibody incubation, tissues were washed with 1X PBS. Alexa-fluor labeled 488 or 555 antibodies were used as secondary antibodies in blocking buffer, and tissues/cells were incubated accordingly with either Hoechst or DAPI for nuclei labeling. Tissues were then washed 1X PBS and mounted. Primary antibodies are found in Table S1.

### Cardiac Tissue Imaging and Actin Alignment Analysis

Confocal images were acquired with a Zeiss LSM 700 Laser Scanning microscope with a 63x (NA 1.4) oil immersion objective to obtain z stacks of the tissue surface. Max intensity projections of z-stacks (n=10 per tissue, about 20 µm depth) were used for cardiomyocyte alignment quantification (actin alignment quantification). A custom MATLAB code was used to quantify the angular distribution of actin filaments, which are orthogonal to the sarcomeres within cardiac tissues based on previous described methods(84). Briefly, a threshold was applied based on actin filament prominence and use to determine which orientations to take into account for the analysis. The α-actinin channel was then used to create a binary mask for cardiomyocyte locations, which was used to determine cardiomyocyte alignment. Using combinations of these masks, we were able to determine the alignment of cardiomyocytes versus fibroblasts within the tissues. Angular distribution of actin filaments was then used to calculate the orientational order parameter (OOP)—a measure of alignment, where OOP values close to 1 indicate completely coaligned actin filaments and values close to 0 indicate an isotropic distribution of actin filaments. An average OOP was obtained for each sample by considering angular distributions of actin filaments within all z-stacks (n=10 per tissue).

### Electron microscopy

Cells/tissues were fixed with 2%glutaraldehyde/2%formaldehyde mixture in 0.1M sodium cacodylate at 4°C. Post-fixation in 1%osmium tetroxide/1.5%potassium ferrocyanide (2 hours at 4°C), cells/tissues were dehydrated using ethanol and embedded in EPON epoxy resin. 60 nm sections were cut transverse in cell direction and contrasted using 5% uranyl acetate in water for 20min followed by Reynolds lead citrate for 2min. Whole nanotomy scans were made of tissues (85).

### RT-PCR

Total RNA was isolated using TRIzol (Sigma). cDNA was synthesized by reverse transcription and real-time PCRs were performed using primers (Table S2) to reveal expression of mRNA transcripts. Relative expression levels were calculated using the ddCt method.

### Statistical Analysis

Data are represented as mean + standard deviation (SD). Statistical analysis was performed using Prism software (GraphPad) or SigmaPlot. Datasets were assessed for normality using a Shapiro-Wilk normality test. Homogeneity of variances was assessed with either Brown-Forsythe or Levene Median Test. Outliers were defined with ROUT (Q=1) testing. Statistical significance was considered a p-value<0.05 and appropriate statistical tests were chosen based on experimental conditions and data. If data did not meet assumptions of normality and homogeneity of variances, data were either log transformed (in order to meet these assumptions) or an appropriate non-parametric test was utilized.

For collagen concentration studies in Supplemental Figure 1, a one-way ANOVA was performed with a post hoc Holm Sidak test. For tissue loading studies shown in Figure 2, 3 and 4, two-way ANOVAs were performed for each dependent variable (diastolic length, diastolic stress, fractional shortening, conduction velocity, cardiomyocyte twitch stress, cardiomyocyte OOP, and conduction velocity) with independent variables being loading condition (constrained or dynamic) and loading amount (1x or 8x). If statistical significance was found via two-way ANOVA, a Holm Sidak post hoc test was performed to determine statistical differences between groups.

For 2D cellular experiments, differences between patient and controls were calculated using an unpaired two-sided *t*-test. Difference between cell lines under either control or 2D stretched condition were calculated with a two-way ANOVA, post-hoc, Bonferroni’s multiple comparisons test. For comparison of elastic moduli of the difference strips (fig. S9), a Mann Whitney U test was performed.

For Figure 6, a two-way ANOVA was performed within each loading amount (either 1x or 8x) for each dependent variable (length, fractional shortening, diastolic stress, contractile force and twitch stress) with independent variables being loading condition (constrained or dynamic) and disease status (control or DSPmut). If statistical significance was found via two-way ANOVA, a Holm Sidak post hoc test was performed to determine statistical differences between groups. **SUPPLEMENTARY MATERIALS**

## ACKNOWLEDGMENTS

Part of the work has been performed at the UMCG Imaging and Microscopy Center (UMIC), for which we express our gratitude. This work was supported by the Human Frontier Science Program [grant number RGY 0071/2014 to P.v.d.M. and A.W.F.], the National Institutes of Health [grant number DP2HL117750 to A.W.F.], the Office of Naval Research [grant number N00014-17-1-2566 to A.W.F.], and the Dowd Fellowship and Presidential Fellowship, Carnegie Mellon University [to J.M.B.].

Carnegie Mellon University has filed for patent protection on the technology described herein, and J.M.B., R.M.D. and A.W.F. are named as inventors on the patent.

## Author contributions

Design study: [A.W.F.] [P.v.d.M.] [J.M.B.] [M.V.]

Establishing methods: [J.M.B.] [M.V.] [M.F.H.] [R.D.] [I.B] [D.S] [J.T] [B.C] [M.C.B]

Conducting experiments: [J.M.B.] [M.V.] [R.D.] [A.K] [A.L] [Y.S] [R.P]

Data processing: [J.M.B.] [M.V.] [D.K.] [I.B]

Data collection: [J.M.B.] [M.V.] [R.P]

Figure contributions: [J.M.B.] [M.V.] [L.V.] [A.S.T.] [I.B] [D.S] [J.T] [R.P] [D.A.P] [A.J.H.S] [J.D.H.J]

Critical data interpretation : ALL

## REFERENCES

1. Lindsey SE, Butcher JT, Yalcin HC. Mechanical regulation of cardiac development. Front Physiol. 2014;5 AUG(August):1–15. doi: 10.3389/fphys.2014.00318

2. Grossman W, Paulus WJ. Myocardial stress and hypertrophy: A complex interface between biophysics and cardiac remodeling. J Clin Invest. 2013. doi: 10.1172/JCI69830

3. Katz AM. Maladaptive growth in the failing heart: The cardiomyopathy of overload. Cardiovasc Drugs Ther. 2002. doi: 10.1023/A:1020604623427

4. Cruz FM, Sanz-Rosa D, Roche-Molina M, et al. Exercise triggers ARVC phenotype in mice expressing a disease-causing mutated version of human plakophilin-2. J Am Coll Cardiol. 2015;65(14):1438–1450. doi: 10.1016/j.jacc.2015.01.045

5. Martherus R, Jain R, Takagi K, et al. Accelerated cardiac remodeling in desmoplakin transgenic mice in response to endurance exercise is associated with perturbed Wnt/β-catenin signaling. Am J Physiol - Hear Circ Physiol. 2015;38103:ajpheart.00295.2015. doi: 10.1152/ajpheart.00295.2015

6. Egido J, Zaragoza C, Gomez-Guerrero C, et al. Animal models of cardiovascular diseases. J Biomed Biotechnol. 2011;2011. doi: 10.1155/2011/497841

7. Feinberg AW, Alford PW, Jin H, et al. Controlling the contractile strength of engineered cardiac muscle by hierarchal tissue architecture. Biomaterials. 2012. doi: 10.1016/j.biomaterials.2012.04.043

8. Dornian IJ, Chiravuri M, Meer VDP, et al. Generation of functional ventricular heart muscle from mouse ventricular progenitor cells. Science (80-). 2009. doi: 10.1126/science.1177350

9. Feinberg AW, Feigel A, Shevkoplyas SS, Sheehy S, Whitesides GM, Parker KK. Muscular thin films for building actuators and powering devices. Science (80-). 2007. doi: 10.1126/science.1146885

10. Feinberg AW, Ripplinger CM, Van Der Meer P, et al. Functional differences in engineered myocardium from embryonic stem cell-derived versus neonatal cardiomyocytes. Stem Cell Reports. 2013. doi: 10.1016/j.stemcr.2013.10.004

11. Wang G, McCain ML, Yang L, et al. Modeling the mitochondrial cardiomyopathy of Barth syndrome with induced pluripotent stem cell and heart-on-chip technologies. Nat Med. 2014. doi: 10.1038/nm.3545

12. Soares CP, Midlej V, Oliveira MEW de, Benchimol M, Costa ML, Mermelstein C. 2D and 3D-Organized Cardiac Cells Shows Differences in Cellular Morphology, Adhesion Junctions, Presence of Myofibrils and Protein Expression. Cordes N, ed. PLoS One. 2012;7(5):e38147. doi: 10.1371/journal.pone.0038147

13. Noorman M, van der Heyden MAG, van Veen TAB, et al. Cardiac cell-cell junctions in health and disease: Electrical versus mechanical coupling. J Mol Cell Cardiol. 2009. doi: 10.1016/j.yjmcc.2009.03.016

14. Naito H, Melnychenko I, Didié M, et al. Optimizing engineered heart tissue for therapeutic applications as surrogate heart muscle. Circulation. 2006. doi: 10.1161/CIRCULATIONAHA.105.001560

15. Hansen A, Eder A, Bönstrup M, et al. Development of a drug screening platform based on engineered heart tissue. Circ Res. 2010. doi: 10.1161/CIRCRESAHA.109.211458

16. Boudou T, Legant WR, Mu A, et al. A microfabricated platform to measure and manipulate the mechanics of engineered cardiac microtissues. Tissue Eng - Part A. 2012. doi: 10.1089/ten.tea.2011.0341

17. Thavandiran N, Dubois N, Mikryukov A, et al. Design and formulation of functional pluripotent stem cell-derived cardiac microtissues. Proc Natl Acad Sci U S A. 2013. doi: 10.1073/pnas.1311120110

18. Zhao Y, Rafatian N, Feric NT, et al. A Platform for Generation of Chamber-Specific Cardiac Tissues and Disease Modeling. Cell. 2019. doi: 10.1016/j.cell.2018.11.042

19. Shadrin IY, Allen BW, Qian Y, et al. Cardiopatch platform enables maturation and scale-up of human pluripotent stem cell-derived engineered heart tissues. Nat Commun. 2017;8(1). doi: 10.1038/s41467-017-01946-x

20. Zhang D, Shadrin IY, Lam J, Xian HQ, Snodgrass HR, Bursac N. Tissue-engineered cardiac patch for advanced functional maturation of human ESC-derived cardiomyocytes. Biomaterials. 2013. doi: 10.1016/j.biomaterials.2013.04.026

21. Li RA, Keung W, Cashman TJ, et al. Bioengineering an electro-mechanically functional miniature ventricular heart chamber from human pluripotent stem cells. Biomaterials. 2018. doi: 10.1016/j.biomaterials.2018.02.024

22. Kupfer ME, Lin W-H, Ravikumar V, et al. In Situ Expansion, Differentiation and Electromechanical Coupling of Human Cardiac Muscle in a 3D Bioprinted, Chambered Organoid. Circ Res. 2020. doi: 10.1161/circresaha.119.316155

23. Ronaldson-Bouchard K, Yeager K, Teles D, et al. Engineering of human cardiac muscle electromechanically matured to an adult-like phenotype. Nat Protoc. 2019. doi: 10.1038/s41596-019-0189-8

24. Ronaldson-Bouchard K, Ma SP, Yeager K, et al. Advanced maturation of human cardiac tissue grown from pluripotent stem cells. Nature. 2018;556(7700):239–243. doi: 10.1038/s41586-018-0016-3

25. Morgan KY, Black LD. Mimicking isovolumic contraction with combined electromechanical stimulation improves the development of engineered cardiac constructs. Tissue Eng - Part A. 2014. doi: 10.1089/ten.tea.2013.0355

26. Ye Morgan K, Black LD. It’s all in the timing Modeling isovolumic contraction through development and disease with a dynamic dual electromechanical bioreactor system. Organogenesis. 2014. doi: 10.4161/org.29207

27. Stoppel WL, Kaplan DL, Black LD. Electrical and Mechanical Stimulation of Cardiac Cells and Tissue Constructs. Vol 96. Elsevier B.V.; 2016:135–155. doi: 10.1016/j.addr.2015.07.009

28. Weinberger F, Mannhardt I, Eschenhagen T. Engineering Cardiac Muscle Tissue: A Maturating Field of Research. Vol 120.; 2017:1487–1500. doi: 10.1161/CIRCRESAHA.117.310738

29. Jallerat Q, Feinberg AW. Extracellular Matrix Structure and Composition in the Early Four-Chambered Embryonic Heart. Cells. 2020. doi: 10.3390/cells9020285

30. Bian W, Jackman CP, Bursac N. Controlling the structural and functional anisotropy of engineered cardiac tissues. Biofabrication. 2014. doi: 10.1088/1758-5082/6/2/024109

31. Liau B, Christoforou N, Leong KW, Bursac N. Pluripotent stem cell-derived cardiac tissue patch with advanced structure and function. Biomaterials. 2011. doi: 10.1016/j.biomaterials.2011.08.050

32. Black LD, Meyers JD, Weinbaum JS, Shvelidze YA, Tranquillo RT. Cell-Induced Alignment Augments Twitch Force in Fibrin Gel-Based Engineered Myocardium via Gap Junction Modification. Tissue Eng - Part A. 2009. doi: 10.1089/ten.tea.2008.0502

33. Blazeski A, Kostecki GM, Tung L. Engineered heart slices for electrophysiological and contractile studies. Biomaterials. 2015. doi: 10.1016/j.biomaterials.2015.03.026

34. Bursac N, Parker KK, Iravanian S, Tung L. Cardiomyocyte cultures with controlled macroscopic anisotropy: a model for functional electrophysiological studies of cardiac muscle. Circ Res. 2002. doi: 10.1161/01.RES.0000047530.88338.EB

35. Kim DH, Lipke EA, Kim P, et al. Nanoscale cues regulate the structure and function of macroscopic cardiac tissue constructs. Proc Natl Acad Sci U S A. 2010. doi: 10.1073/pnas.0906504107

36. Thomas SP, Bircher-Lehmann L, Thomas SA, Zhuang J, Saffitz JE, Kléber AG. Synthetic strands of neonatal mouse cardiac myocytes: Structural and electrophysiological properties. Circ Res. 2000. doi: 10.1161/01.RES.87.6.467

37. Horváth A, Lemoine MD, Löser A, et al. Low Resting Membrane Potential and Low Inward Rectifier Potassium Currents Are Not Inherent Features of hiPSC-Derived Cardiomyocytes. Stem Cell Reports. 2018. doi: 10.1016/j.stemcr.2018.01.012

38. Halbach M, Pillekamp F, Brockmeier K, Hescheler J, Müller-Ehmsen J, Reppel M. Ventricular slices of adult mouse hearts - A new multicellular in vitro model for electrophysiological studies. Cell Physiol Biochem. 2006. doi: 10.1159/000095132

39. Santana LF, Cheng EP, Lederer WJ. How does the shape of the cardiac action potential control calcium signaling and contraction in the heart? J Mol Cell Cardiol. 2010. doi: 10.1016/j.yjmcc.2010.09.005

40. Nunes SS, Miklas JW, Liu J, et al. Biowire: A platform for maturation of human pluripotent stem cell-derived cardiomyocytes. Nat Methods. 2013. doi: 10.1038/nmeth.2524

41. Maron BJ, Chaitman BR, Ackerman MJ, et al. Recommendations for physical activity and recreational sports participation for young patients with genetic cardiovascular diseases. Circulation. 2004. doi: 10.1161/01.CIR.0000128363.85581.E1

42. Kirchhof P, Fabritz L, Zwiener M, et al. Age- and training-dependent development of arrhythmogenic right ventricular cardiomyopathy in heterozygous plakoglobin-deficient mice. Circulation. 2006. doi: 10.1161/CIRCULATIONAHA.106.624502

43. Moncayo-Arlandi J, Guasch E, la Garza MS de, et al. Molecular disturbance underlies to arrhythmogenic cardiomyopathy induced by transgene content, age and exercise in a truncated PKP2 mouse model. Hum Mol Genet. 2016. doi: 10.1093/hmg/ddw213

44. Lyon RC, Mezzano V, Wright AT, et al. Connexin defects underlie arrhythmogenic right ventricular cardiomyopathy in a novel mouse model. Hum Mol Genet. 2014. doi: 10.1093/hmg/ddt508

45. te Rijdt WP, van der Klooster ZJ, Hoorntje ET, et al. Phospholamban immunostaining is a highly sensitive and specific method for diagnosing phospholamban p.Arg14del cardiomyopathy. Cardiovasc Pathol. 2017. doi: 10.1016/j.carpath.2017.05.004

46. Hinson JT, Chopra A, Nafissi N, et al. Titin mutations in iPS cells define sarcomere insufficiency as a cause of dilated cardiomyopathy. Science (80-). 2015;349(6251):982–986. doi: 10.1126/science.aaa5458

47. Ma Z, Huebsch N, Koo S, et al. Contractile deficits in engineered cardiac microtissues as a result of MYBPC3 deficiency and mechanical overload. Nat Biomed Eng. 2018. doi: 10.1038/s41551-018-0280-4

48. Hershberger RE, Hedges DJ, Morales A. Dilated cardiomyopathy: The complexity of a diverse genetic architecture. Nat Rev Cardiol. 2013. doi: 10.1038/nrcardio.2013.105

49. Chattergoon NN, Giraud GD, Louey S, Stork P, Fowden AL, Thornburg KL. Thyroid hormone drives fetal cardiomyocyte maturation. FASEB J. 2012;26(1):397–408. doi: 10.1096/fj.10-179895

50. Parikh SS, Blackwell DJ, Gomez-Hurtado N, et al. Thyroid and Glucocorticoid Hormones Promote Functional T-Tubule Development in Human-Induced Pluripotent Stem Cell-Derived Cardiomyocytes. Circ Res. 2017;121(12):1323–1330. doi: 10.1161/CIRCRESAHA.117.311920

51. Burchfield JS, Xie M, Hill JA. Pathological ventricular remodeling: Mechanisms: Part 1 of 2. Circulation. 2013. doi: 10.1161/CIRCULATIONAHA.113.001878

52. Voorhees AP, Han HC. Biomechanics of cardiac function. Compr Physiol. 2015. doi: 10.1002/cphy.c140070

53. Moustakidis P, Maniar HS, Cupps BP, et al. Altered left ventricular geometry changes the border zone temporal distribution of stress in an experimental model of left ventricular aneurysm: A finite element model study. Circulation. 2002. doi: 10.1161/01.cir.0000032898.55215.0d

54. Sáez P, Kuhl E. Computational modeling of acute myocardial infarction. Comput Methods Biomech Biomed Engin. 2016. doi: 10.1080/10255842.2015.1105965

55. Hayashida W, Kumada T, Nohara R, et al. Left ventricular regional wall stress in dilated cardiomyopathy. Circulation. 1990. doi: 10.1161/01.CIR.82.6.2075

56. Scardulla F, Rinaudo A, Pasta S, Scardulla C. Evaluation of ventricular wall stress and cardiac function in patients with dilated cardiomyopathy. Proc Inst Mech Eng Part H J Eng Med. 2016. doi: 10.1177/0954411915617984

57. Di Napoli P, Taccardi AA, Grilli A, et al. Left ventricular wall stress as a direct correlate of cardiomyocyte apoptosis in patients with severe dilated cardiomyopathy. Am Heart J. 2003. doi: 10.1016/S0002-8703(03)00445-9

58. D’Amore A, Yoshizumi T, Luketich SK, et al. Bi-layered polyurethane – Extracellular matrix cardiac patch improves ischemic ventricular wall remodeling in a rat model. Biomaterials. 2016. doi: 10.1016/j.biomaterials.2016.07.039

59. Matsumura Y, Zhu Y, Jiang H, et al. Intramyocardial injection of a fully synthetic hydrogel attenuates left ventricular remodeling post myocardial infarction. Biomaterials. 2019. doi: 10.1016/j.biomaterials.2019.119289

60. Lee A, Hudson AR, Shiwarski DJ, et al. 3D bioprinting of collagen to rebuild components of the human heart. Science (80-). 2019. doi: 10.1126/science.aav9051

61. Macqueen LA, Sheehy SP, Chantre CO, et al. A tissue-engineered scale model of the heart ventricle. Nat Biomed Eng. 2018. doi: 10.1038/s41551-018-0271-5

62. Keung W, Chan PKW, Backeris PC, et al. Human Cardiac Ventricular-Like Organoid Chambers and Tissue Strips From Pluripotent Stem Cells as a Two-Tiered Assay for Inotropic Responses. Clin Pharmacol Ther. 2019. doi: 10.1002/cpt.1385

63. Cokkinos D V., Belogianneas C. Left ventricular remodelling: A problem in search of solutions. Eur Cardiol Rev. 2016. doi: 10.15420/ecr.2015:9:3

64. Sabbah HN. Silent disease progression in clinically stable heart failure. Eur J Heart Fail. 2017. doi: 10.1002/ejhf.705

65. Seidman JG, Seidman C. The Genetic Basis for Cardiomyopathy. Cell. 2001. doi: 10.1016/s0092-8674(01)00242-2

66. St. John Sutton MG, Sharpe N. Left ventricular remodeling after myocardial infarction: Pathophysiology and therapy. Circulation. 2000. doi: 10.1161/01.cir.101.25.2981

67. Parvari R, Levitas A. The mutations associated with dilated cardiomyopathy. Biochem Res Int. 2012. doi: 10.1155/2012/639250

68. Dellefave L, McNally EM. The genetics of dilated cardiomyopathy. Curr Opin Cardiol. 2010. doi: 10.1097/HCO.0b013e328337ba52

69. McNally E, Allikian M, Wheeler MT, Mislow JM, Heydemann A. Cytoskeletal defects in cardiomyopathy. J Mol Cell Cardiol. 2003. doi: 10.1016/S0022-2828(03)00018-X

70. Sheikh F, Ross RS, Chen J. Cell-Cell Connection to Cardiac Disease. Trends Cardiovasc Med. 2009. doi: 10.1016/j.tcm.2009.12.001

71. Buck VU, Hodecker M, Eisner S, Leube RE, Krusche CA, Classen-Linke I. Ultrastructural changes in endometrial desmosomes of desmoglein 2 mutant mice. Cell Tissue Res. 2018;374(2):317–327. doi: 10.1007/s00441-018-2869-z

72. Travers JG, Kamal FA, Robbins J, Yutzey KE, Blaxall BC. Cardiac fibrosis: The fibroblast awakens. Circ Res. 2016. doi: 10.1161/CIRCRESAHA.115.306565

73. Zhao X, Tan RS, Tang HC, et al. Left ventricular wall stress is sensitive marker of hypertrophic cardiomyopathy with preserved ejection fraction. Front Physiol. 2018. doi: 10.3389/fphys.2018.00250

74. Montgomery M, Ahadian S, Davenport Huyer L, et al. Flexible shape-memory scaffold for minimally invasive delivery of functional tissues. Nat Mater. 2017. doi: 10.1038/nmat4956

75. Weinberger F, Breckwoldt K, Pecha S, et al. Cardiac repair in Guinea pigs with human engineered heart tissue from induced pluripotent stem cells. Sci Transl Med. 2016. doi: 10.1126/scitranslmed.aaf8781

76. Ovchinnikova E, Hoes M, Ustyantsev K, et al. Modeling Human Cardiac Hypertrophy in Stem Cell-Derived Cardiomyocytes. Stem Cell Reports. 2018;10(3):794–807. doi: 10.1016/j.stemcr.2018.01.016

77. Burridge PW, Matsa E, Shukla P, et al. Chemically defned generation of human cardiomyocytes. Nat Methods. 2014;11(8):855–860. doi: 10.1038/nMeth.2999

78. Lian X, Zhang J, Azarin SM, et al. Directed cardiomyocyte differentiation from human pluripotent stem cells by modulating Wnt/??-catenin signaling under fully defined conditions. Nat Protoc. 2013;8(1):162–175. doi: 10.1038/nprot.2012.150

79. Xiu QX, Set YS, Sun W, Zweigerdt R. Global expression profile of highly enriched cardiomyocytes derived from human embryonic stem cells. Stem Cells. 2009;27(9):2163–2174. doi: 10.1002/stem.166

80. Okita K, Matsumura Y, Sato Y, et al. A more efficient method to generate integration-free human iPS cells. Nat Methods. 2011;8(5):409–412. doi: 10.1038/nmeth.1591

81. Takahashi K, Tanabe K, Ohnuki M, et al. Induction of Pluripotent Stem Cells from Adult Human Fibroblasts by Defined Factors. Cell. 2007;131(5):861–872. doi: 10.1016/j.cell.2007.11.019

82. Tohyama S, Hattori F, Sano M, et al. Distinct metabolic flow enables large-scale purification of mouse and human pluripotent stem cell-derived cardiomyocytes. Cell Stem Cell. 2013;12(1):127–137. doi: 10.1016/j.stem.2012.09.013

83. Martin L, Grigoryan A, Wang D, et al. Identification and characterization of small molecules that inhibit nonsense-mediated rna decay and suppress nonsense p53 mutations. Cancer Res. 2014;74(11):3104–3113. doi: 10.1158/0008-5472.CAN-13-2235

84. Sun Y, Jallerat Q, Szymanski JM, Feinberg AW. Conformal nanopatterning of extracellular matrix proteins onto topographically complex surfaces. Nat Methods. 2015;12(2):134–136. doi: 10.1038/nmeth.3210

85. Sokol E, Kramer D, Diercks GFH, et al. Large-scale electron microscopy maps of patient skin and mucosa provide insight into pathogenesis of blistering diseases. J Invest Dermatol. 2015. doi: 10.1038/jid.2015.109

