## Supplemental Methods, Figures, and Tables for "Dynamic Loading of Human Engineered Heart Tissue Enhances Contractile Function and Drives Desmosome-linked Disease Phenotype"

**SUPPLEMENTARY METHODS**

**Validation of Strip Bending Force Measurements**

Finite element modelling of forces associated with strip bending (fig. S3A) were validated by measuring the forces associated with strip bending using a custom-mounted optical force transducer (World Precision Instruments,  SI-KG20). Linear actuators moving the force transducer were used to bend the strip and measure the forces associated with strip bending (Figure S3). The linear actuators were moved to discrete tissue lengths (4.5, 7.5, and 9.5 mm) and then compared to the model. Both x and z deformation forces were obtained with strip modeling; however, the z component of bending was orders of magnitude lower than the x component (defined in Figure S3) of bending in both 1x and 8x strips (fig. S3A). Thus, only the x component of bending was used to validate the strip bending forces (fig. S3B). A schematic of the force transducer set-up utilized to assess strip x-bending is shown (fig. S3C). Representative force transducer measurements obtained from x bending of strip over lengths associated with normal tissue deformations were in good agreement with those observed from finite element modeling of 8x strips as demonstrated in Figure 1.

**Inter-desmosomal space estimation**

Images (2D cultured cells) of desmosomes and cell-cell borders were taken with a CM100 TEM microscope at a 46.000x magnification. ImageJ was used to make intensity plots of up to 50 desmosomes per condition. To determine the inter-desmosomal space, the length between the two highest intensity peaks (located approximately within both membranes/outer dense plaques) was measured.

**SUPPLEMENTARY FIGURES AND FIGURE LEGENDS**

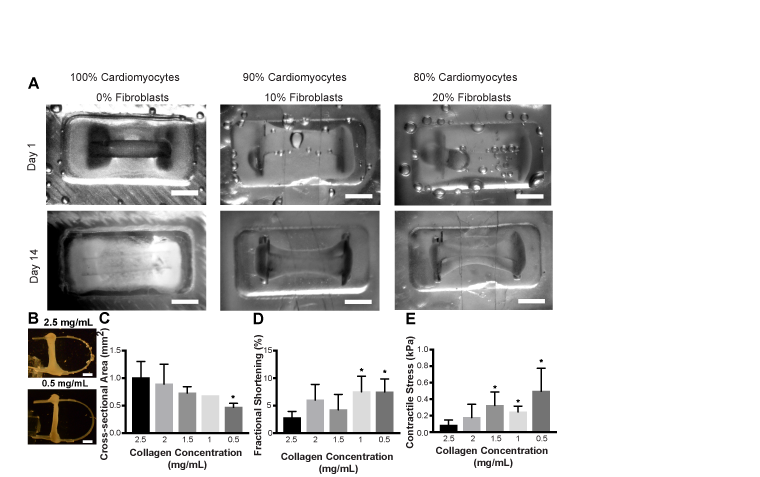

**Fig S1. Impact of Collagen and Fibroblast Concentration on Tissue Morphology and Function.** (A) Macroscopic images during tissue compaction with different ratios of cardiomyocytes to cardiac fibroblasts. (B) Tissue macroscopic images from the side showing differences in tissue cross-sectional area when different casting collagen concentrations are used. Tissues were made with 10% fibroblasts. Scale bars are 2 mm. (C) Average tissue cross-sectional area with collagen concentrations ranging from 0.5 mg/mL to 2.5 mg/mL. (D) Average tissue contractile strain in tissues fabricated with 0.5 mg/mL to 2.5 mg/mL collagen. (E) Cardiomyocyte generated twitch stress in tissues fabricated with different collagen concentrations (0.5 mg/mL to 2.5 mg/mL). Results are based on one-way ANOVA with post hoc Holm Sidak test. * indicates p<0.05 compared to 2.5 mg/mL collagen.

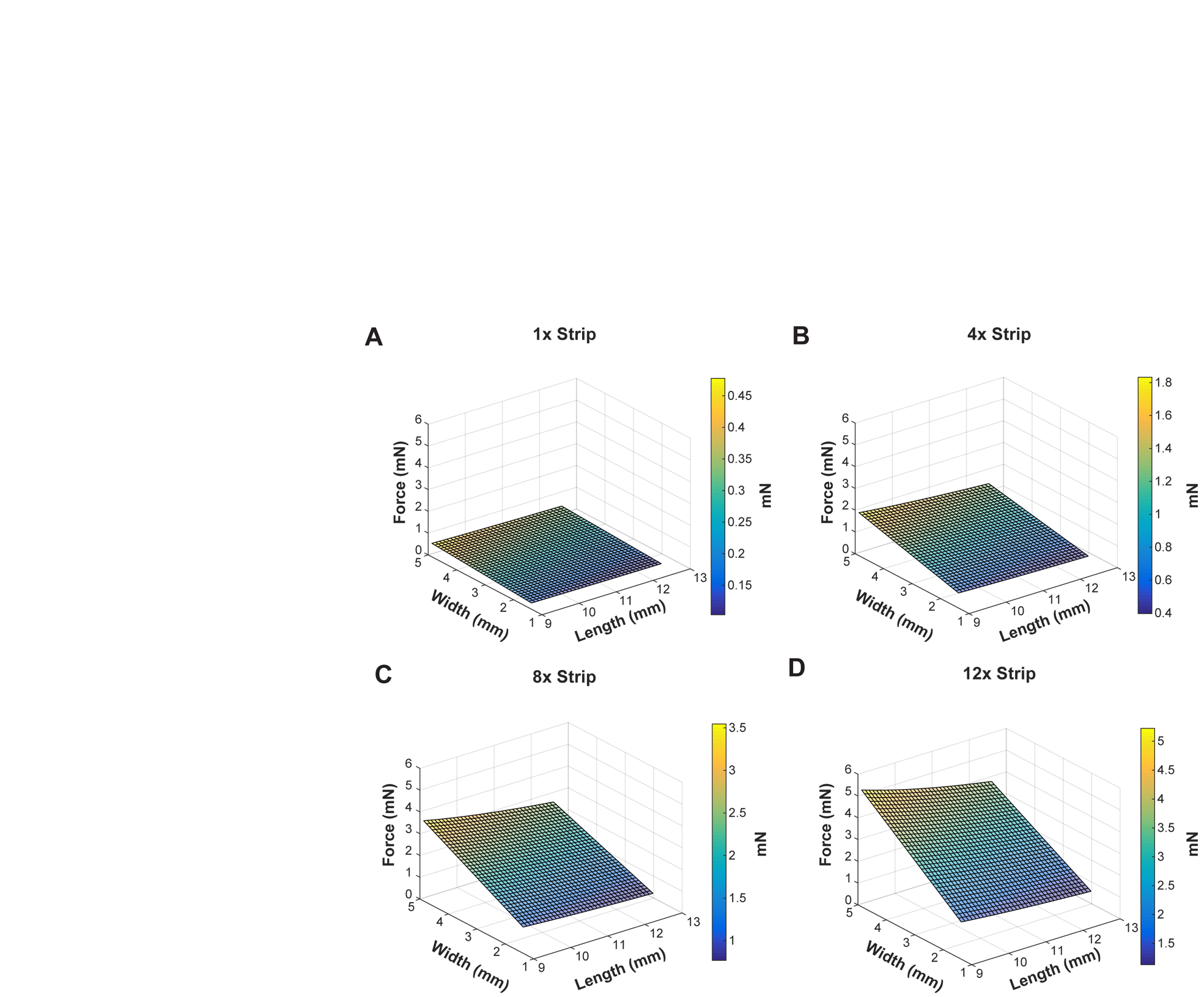

**Fig. S2. Altering Strip Dimensions for a Specified Force Output.** Surface plot of forces associated with different strip widths and length of the strip between tissue attachment points. Tissue length was kept constant (6 mm). Graphs are shown for (A) 1x strips (B) 4x strips (C) 8x Strips and (D) 12x strips.

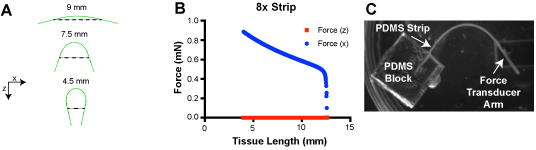
**Fig S3. Force Transducer Validation of Strip Bending Forces. (**A) Strip deformation associated with different tissue lengths (indicated by dotted lines). X and z displacements were applied to the strip at tissue attachment points. (B) Respective contributions of x and z bending reaction forces with strip deformation during modeling observed in 8x strips. (C) Force Transducer set-up used to validate force measurements obtained from strips. All measurements were obtained via x bending of 8x strip.

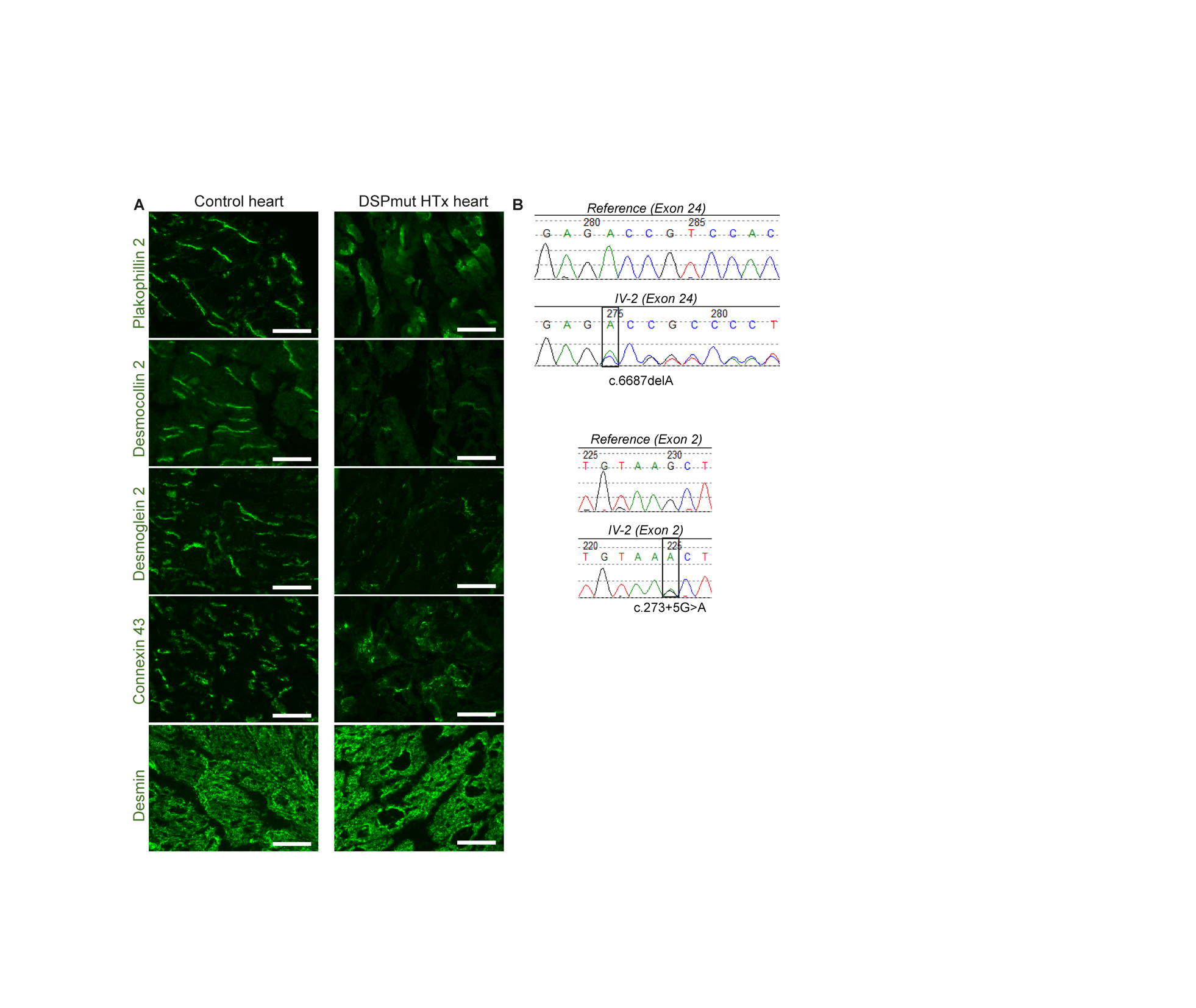

**Fig. S4. *DSP* Mutations and desmosomal staining on DSPmut patient heart.** (A) Sanger sequencing of both *DSP* alleles of DSPmut patient (IV-2) showing both c.273+5G>A and c.6687delA mutations. (B) Panel indicates immunofluorescent (IF) staining of desmosomal proteins, Cx43 and desmin in human control and heart tissue of DSPmut patient (DSPmut HTx Heart). Scale bars are 50 µm.

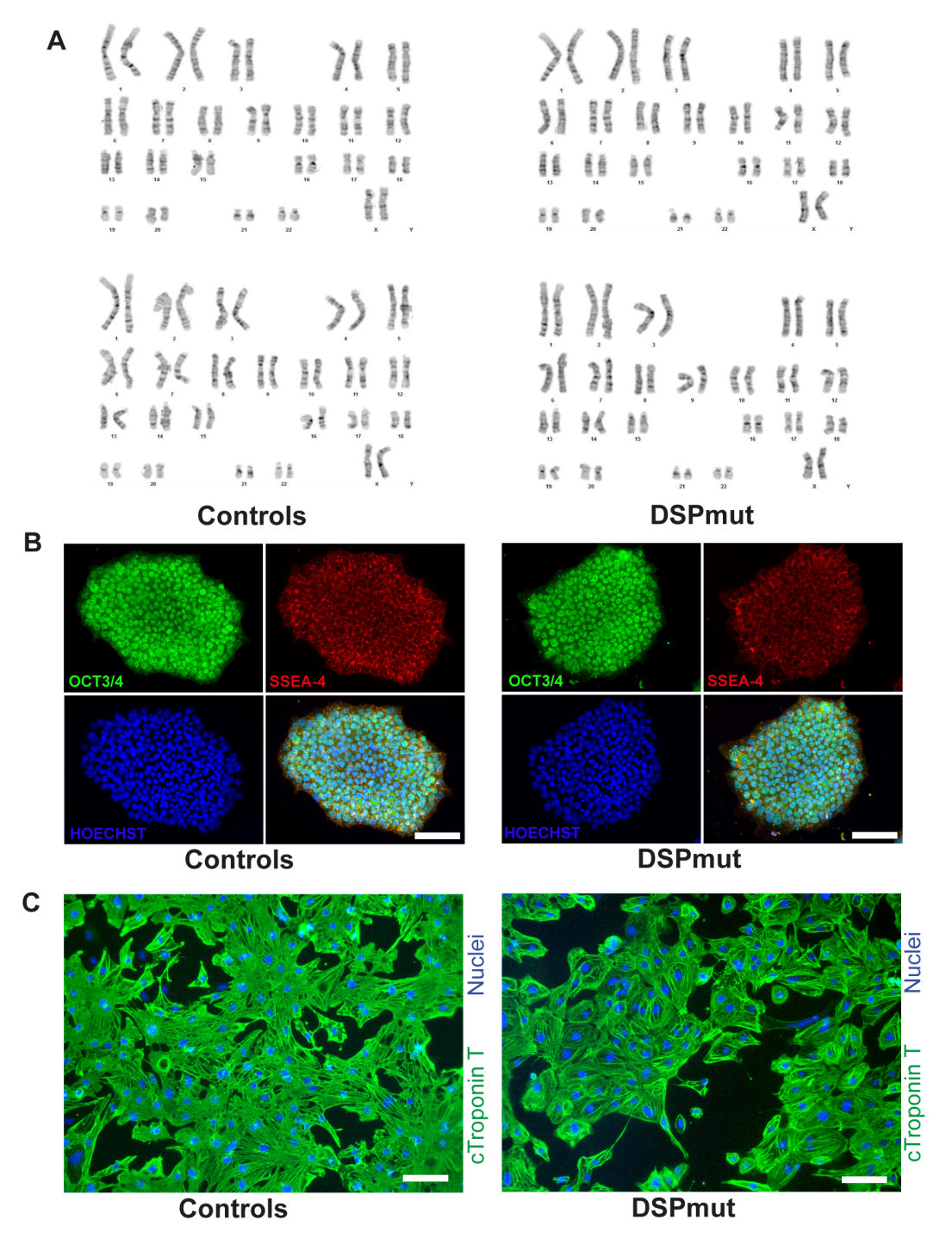

**Fig. S5. Validation of hiPSC lines.** (A) IF staining of pluripotency markers OCT3/4 and SSEA-4 of hiPSC colonies after passage 10. Scale bars are 100 µm. (B) Karyotypes of all hiPSC lines used in this study. (C) IF staining of cardiac troponin T on hiPSC-differentiated cardiomyocytes. Scale bars are 50 µm.

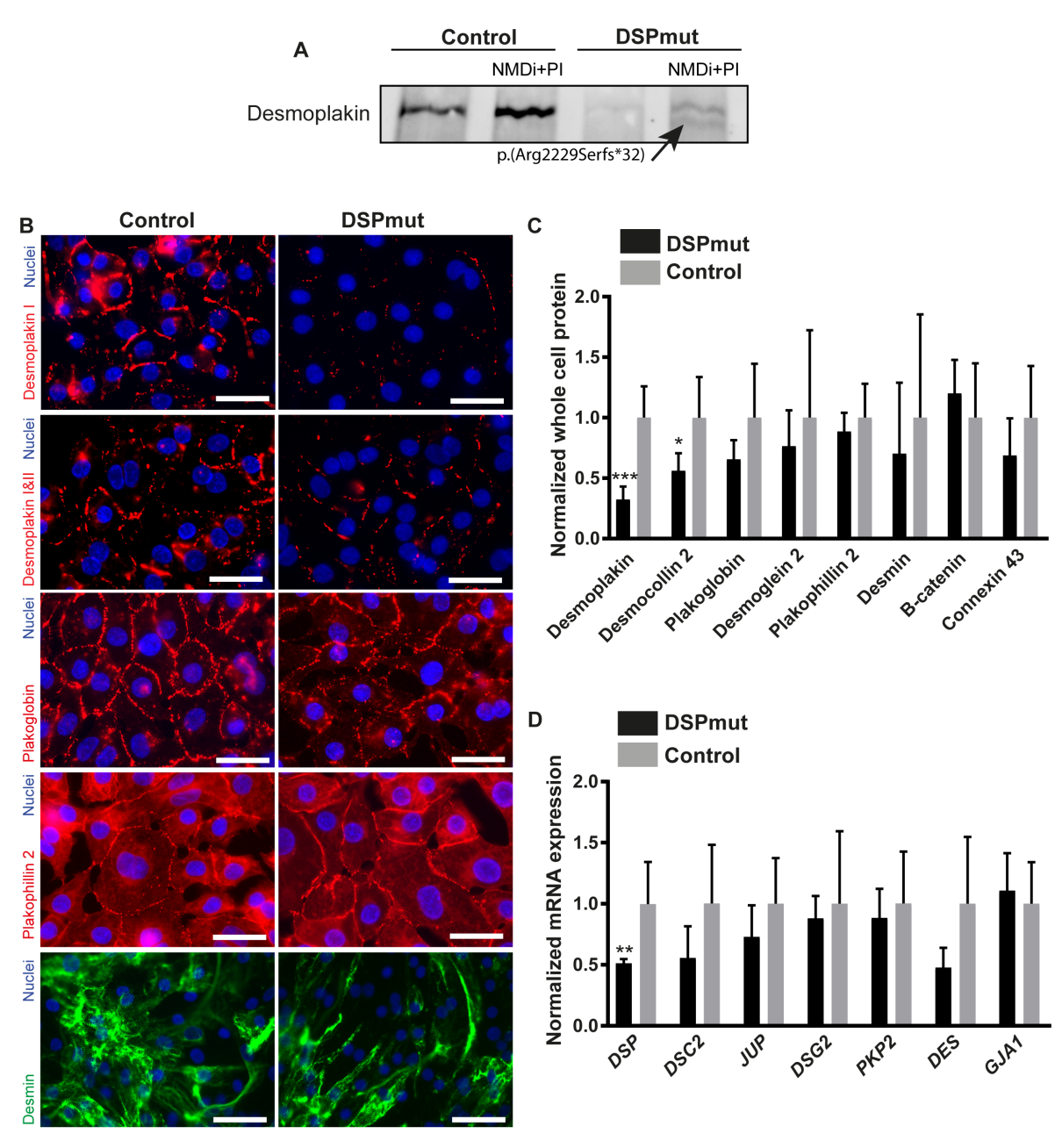

**Fig. S6. Baseline levels of desmosomal proteins and genes cultured in 2D**. (A) Nonsense mediated decay is responsible for breakdown of desmoplakin product p.(Arg2229Serfs*32) originating from the DSP allele with the c.6687delA mutation. Blot shows desmoplakin expression in control and DSPmut cells under nonsense mediated decay inhibitor (NMDi) and proteasome inhibitor (PI, bortezomib). (B) IF staining of key desmosomal proteins and desmin in cardiomyocytes at baseline. Scale bars are 50 µm. (C) Fold change expression of desmosomal proteins, beta-catenin and Cx43 measured in cardiomyocytes determined by western blot. n=6 vs. 4 (patient vs. control); * p<0.05 (unpaired two-sided t-test compared to control);*** p<0.05 (unpaired two-sided t-test compared to control). (D) mRNA expression levels of desmosomal genes. n=5; ** p<0.05 (unpaired two-sided t-test compared to control).

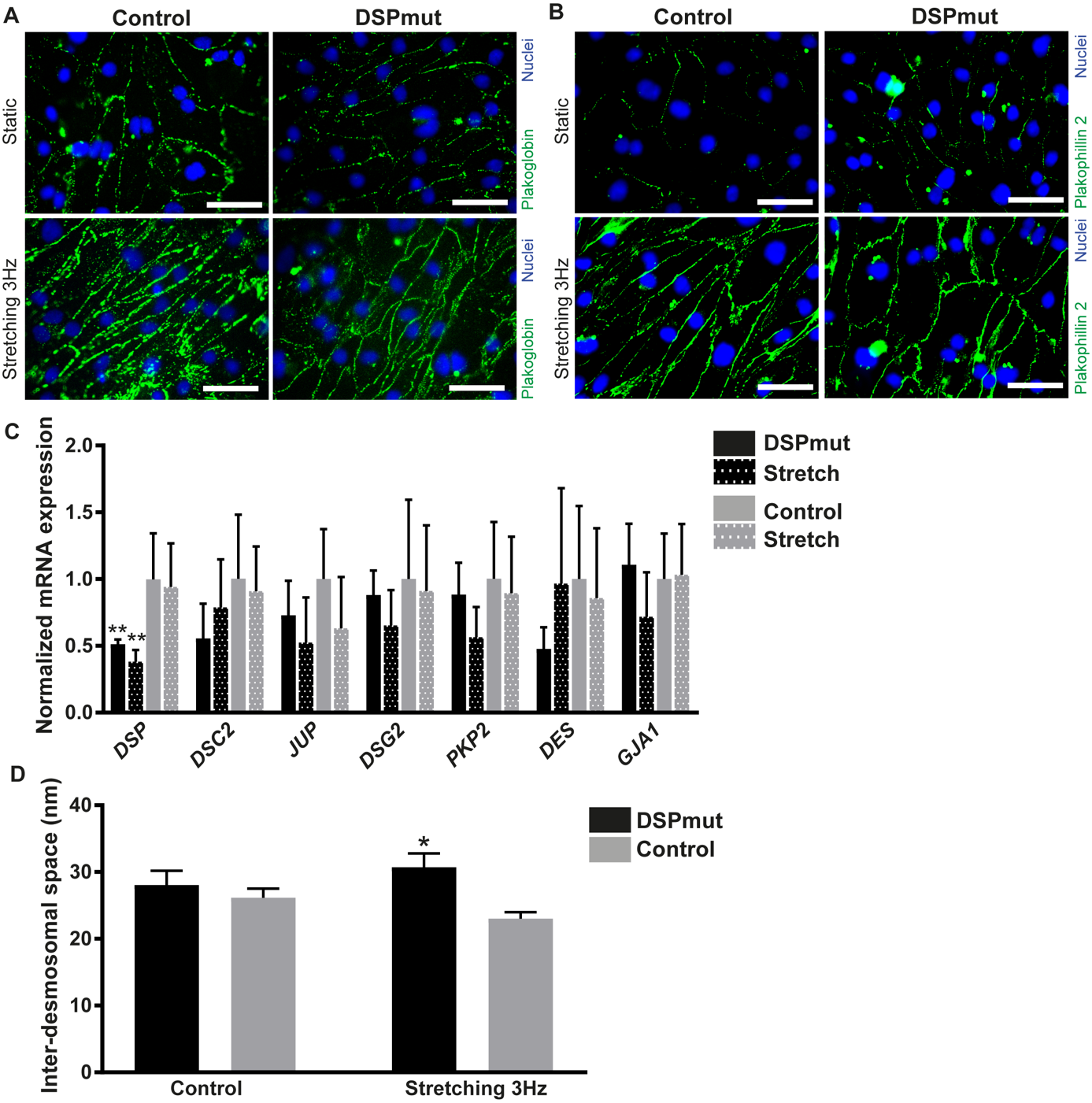

**Fig. S7. Desmosomal protein and gene expression with 2D stretch**. (A-B) IF staining of PG, PKP2 in cardiomyocytes at baseline and after 3Hz stretch. Scale bars are 50 µm. (C) mRNA expression levels of desmosomal genes at baseline and after 3Hz stretch. ** p<0.05 (two-way ANOVA, post hoc Bonferroni’s multiple comparisons test). (D) A proxy of inter-desmosomal space was measured based on 50 desmosomes in each group at baseline and after 3Hz stretch. * p<0.05 (two-way ANOVA, post hoc Bonferroni’s multiple comparisons test).

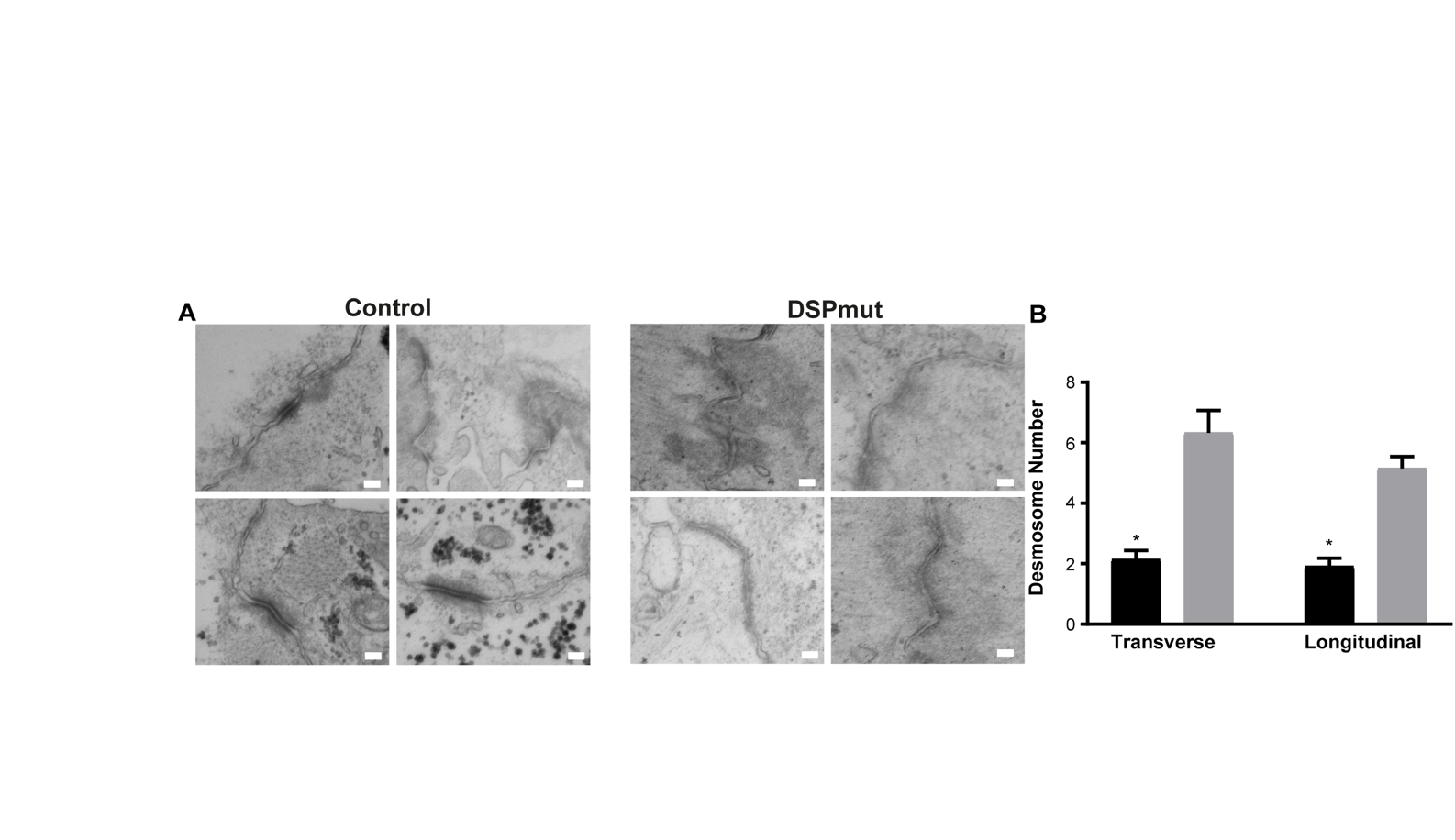
**Fig. S8. Electron microscopy overview in 3D.** (A) Panels depict representative overview pictures of intercalated disc found throughout patient and control 3D tissues. Scale bars are 150 nm. . (B) Desmosomal number per TEM high powered field in control and DSPmut 8x dyn-EHTs. *p<0.05 unpaired t-test compared to control.

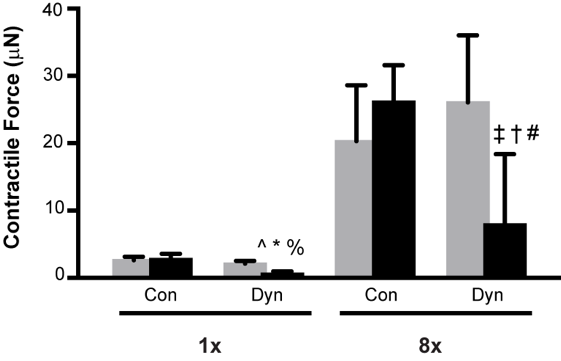

**Fig. S9. Additional Disease Modelling Data.** (A) Average tissue contractile force at day 28 in control and DSPmut tissues under all loading conditions. Results are based on two-way ANOVA with post hoc Holm Sidak test. ^ p<0.05 versus Control 1x constrained. * p<0.05 versus Control 1x dynamic. % p<0.05 versus DSPmut 1x constrained. ‡ p<0.05 control 8x constrained. † p<0.05 from control 8x dynamic. # p<0.05 from DSPmut 8x constrained .

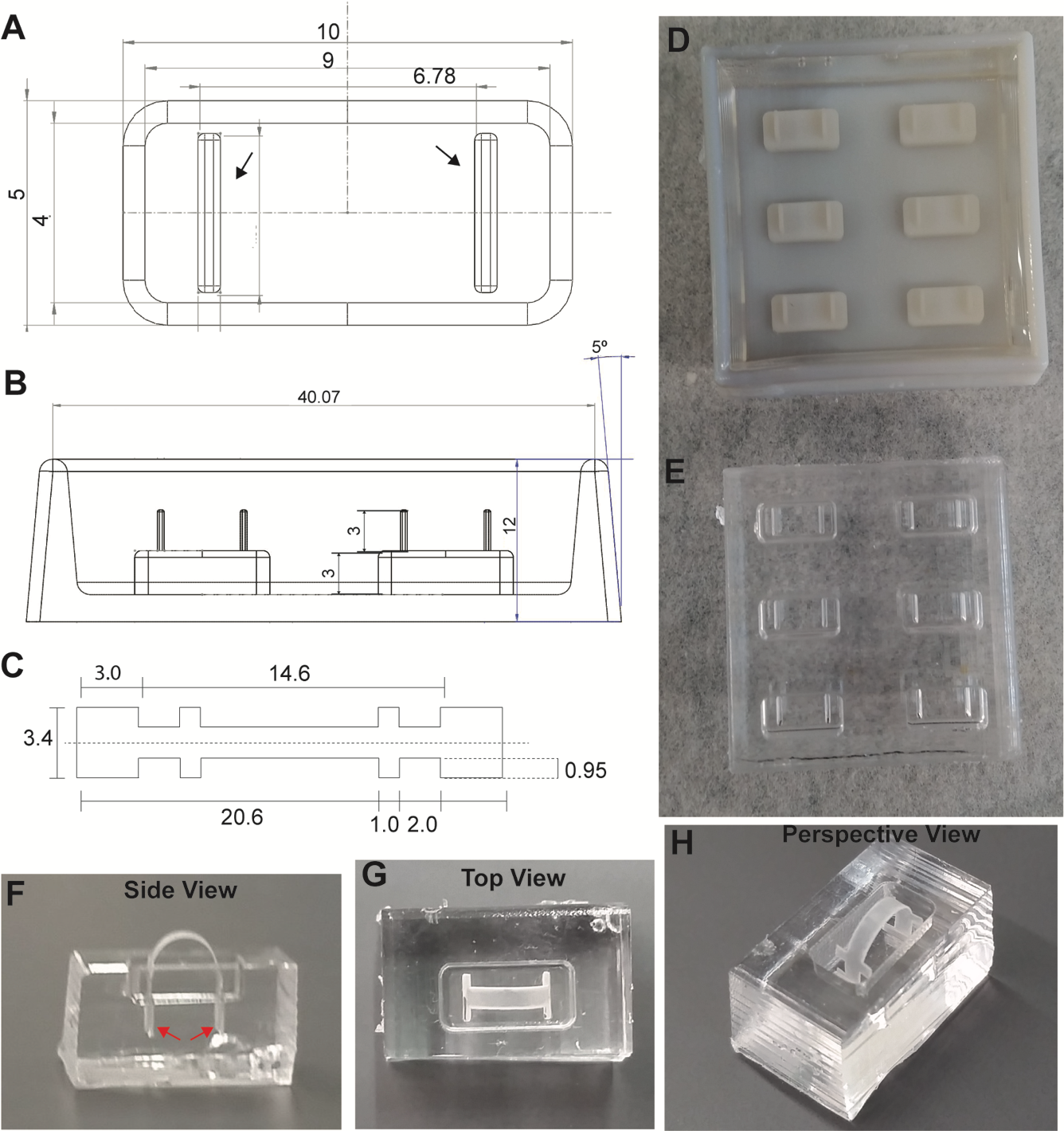

**Fig. S10.** **Mold and Strip Design and Fabrication**. (A) Aerial view of plastic mold dimensions used to create individual PDMS wells for tissue casting. Raised portions of the plastic molds (indicated by the arrows) served as slots in the casted well where the PDMS Strip would eventually sit. (B) Side view of dimensions for plastic mold used to create individual PDMS wells. (C) PDMS strip dimensions demonstrating a 1 mm wide necking region for tissue attachment and a small 3.4 mm wide overhang to prevent tissue from sliding up the PDMS strip during culture and contraction studies. (D) Aerial image of plastic mold used for casting to create PDMS wells. (E) PDMS was cast into wells to create multiple PDMS wells which could be cut out to form individual PDMS wells for tissue fabrication. (F-G) Side, Top and Perspective Views of the PDMS strip placed in the PDMS well design. The PDMS strip sits within the slits in the bottom of the PDMS well (red arrows). All dimensions are in mm.

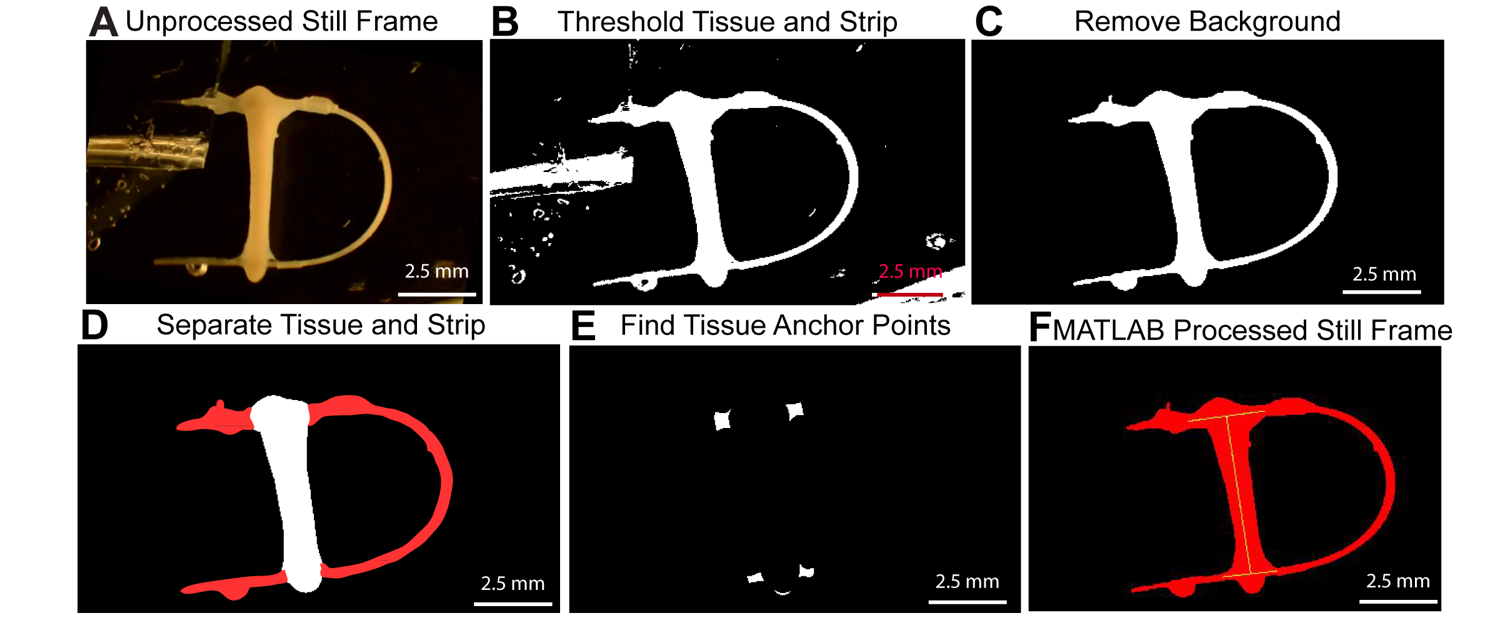
**Fig. S11. MATLAB Processing of Tissue Contractility Videos.** (A) Tissue contraction videos are taken on a stereomicroscope equipped with a Nikon DSLR camera. A representative still frame of the tissue is shown. (B) Tissue videos are uploaded into a MATLAB program where a user threshold the tissue and the strip. The thresholded region is shown in white. (C) Thresholded area that is neither the tissue nor the strip, also known as background, is removed. (D) The program then separates the tissue from its associated strip. (E) Tissue anchor points, or where the tissue attaches to the strip, is then detected. This is used to calculate the length between tissue attachment points on the strip. (F) Tissue length can then be determined based on the change in distance between tissue attachment points over time. Scale bars are 2.5 mm.

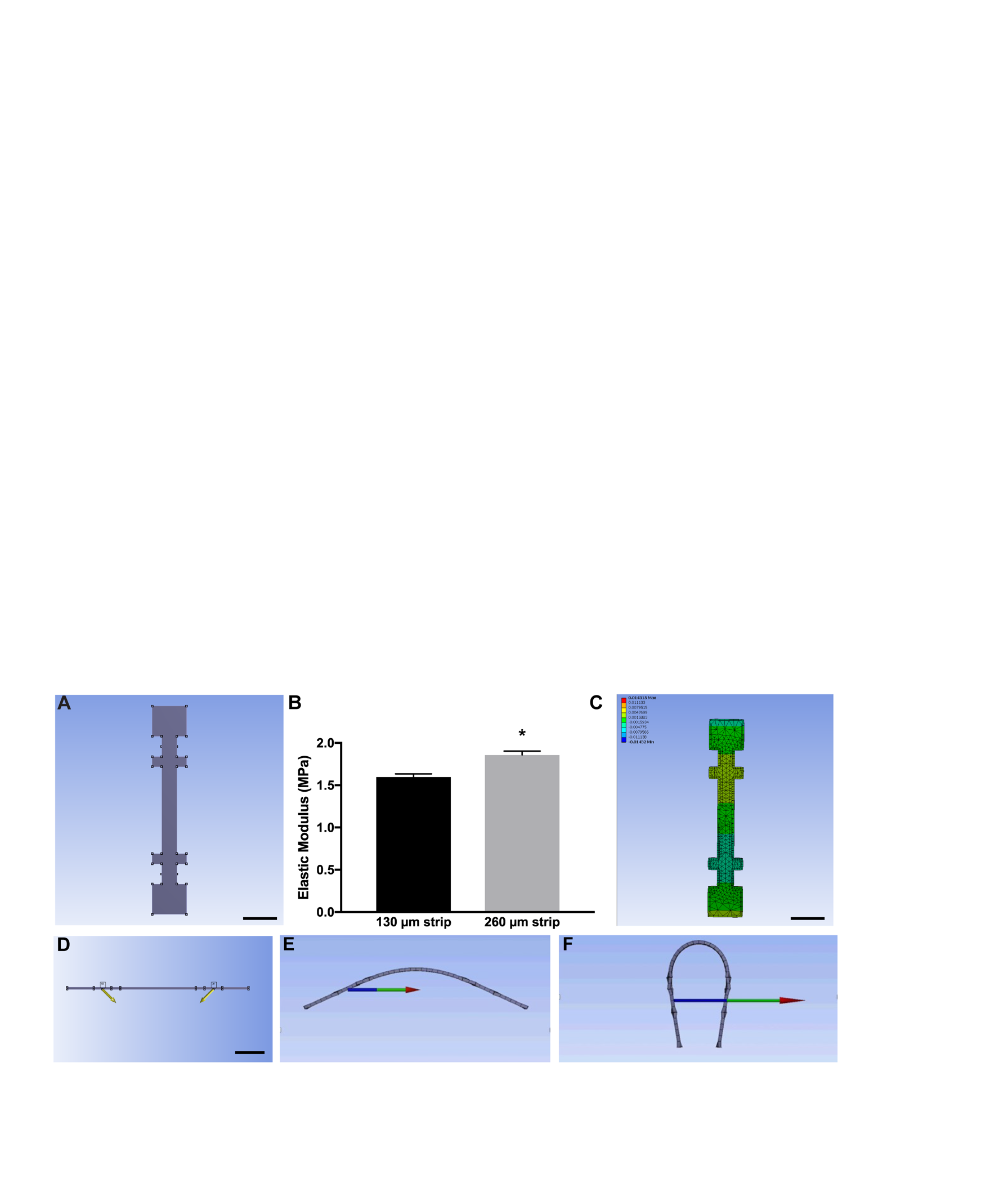
**Fig. S12. Finite Element Modeling (FEM) of Force Required to Bend PDMS Strip.** (A) A 3D model of strip was created using ANSYS SpaceClaim software. (B) Material properties (elastic modulus and Poisson’s ratio) and assumptions were entered in order to model material deformation. Bar graph showing average elastic modulus of each strip used in these studies. * p=0.0043 (Mann-Whitney U test on ranks). (C) Program generated mesh used for modeling. (D) The PDMS strip was modeled as initially flat and displacements (indicated by the yellow arrows) were applied to the tissue attachments points. Reaction forces (i.e. force required to deform the strip under such a displacement) were obtained and examples of relative reaction forces (colored arrows) are shown comparing minimal strip bending (E) and large strip bending (F). Relative increase in length of arrow indicates the increase in force required to bend the strip at larger deformations.

| Table S1:Immunolabeling |  |  |  |
| --- | --- | --- | --- |
| Protein | **Clone/ Cat no.** | **Clonality** | **Usage** |
| Desmoplakin-I (ROD) | DP2.17 | moAb | WB, IHC-Fr, IF |
| Desmoplakin-I&II (C-ter) | 2A5 | moAb | WB, IHC-Fr, IF |
| Plakoglobin | 15F11 | moAb | WB, IHC-Fr, IF |
| Plakophilin 2 | PP2/62;8/86;2/150 | moAb | WB, IHC-Fr, IF |
| Desmoglein 2 | 10G11 | moAb | WB, IHC-Fr |
| Desmocollin 2 | 610120 (Progen) | poAb | WB, IHC-Fr |
| Desmin | Y-20 (Santa Cruz) | poAb | WB, IF |
| Desmin | DE-R-11 | moAb | ICH-Fr |
| Connexin 43 | H-150 (Santa Cruz) | poAb | IHC-Fr |
| cardiac Troponin T | ab45932 (Abcam) | poAb | IF |
| GAPDH | 10R-G109a | moAb | WB |
| Vinculin | SPM227 | moAb | WB |
| SSEA-4 | MC813 | moAb | IF |
| OCT3/4 | H-134 | moAb | IF |
| Alexa Fluor™ 488 Phalloidin | A12379 (Invitrogen) | NA | IF |

| Table S2: RT-PCR | |  |  |
| --- | --- | --- | --- |
| Gene |  | **Forward '5 - '3** | **Reverse '3 - '5** |
| *DSP (NG_008803.1)* |  | CAGTGGTGTCAGCGATGATGT | TGACGCTGGATATGGTGGAA |
|  | *DSP_ex1-4* | GGACGGCTACTG**TCAAACC** | GACTCGAGGGACACTGATG |
|  | *DSP_ex2-5* | **CT**GTCAAACCGGCACGATGTC | TCCAGCTGCCAGCGATAGTC |
|  | *DSP_ex2-int2* | AGGCACCAGAACCAGAACAC | CCCAACCCAGGAACAGAAAC |
|  | *DSP_ex1-int2A* | GGACGGCTACTG**TCAAACC** | CCCAACCCAGGAACAGAAAC |
| *JUP* |  | AGTAGCCACGATGGAGGTGA | AGGTGTATGTCTGCTGCCAC |
| *PKP2* |  | GCAAATGGTTTGCTCGATTT | GGCTGGTAATCTGCAATGGT |
| *DSG2* |  | TCCACTATGCCACCAACCAC | GCTGGAGCATACACCCTCTC |
| *DSC2* |  | CGGAGATTGTTGCGGTTGA | GGAAAGACGTGCTGCTGTATCA |
| *DES* |  | CTGAGCAAAGGGGTTCTGAG | ACTTCATGCTGCTGCTGTGT |
| *GJA1* |  | GGAATGCAAGAGAGGTTGAAAG | GGCATTTGGAGAAACTGGTAGA |
| *PPIA* |  | ACTTCACACGCCATAATG | ACCCGTATGCTTTAGGAT |
| *MYH6* |  | GATAGAGAGACTCCTGCGGC | TCGGTCATCTTGGTGCTTCC |
| *MYH7* |  | CGAAGGGCTTGAATGAGGAGT | TCCTCCCAAGGAGCTGTTAC |
| *TTN* | N2B | CACTAACTGTGACAGTGCC | CTTCTTCCTTTGGTTCAGGT |
| *TTN* | N2A | ATGGAAATGAAAGCTGCCC | GGTGAATTTGGCTAGGTGG |
| *TNNI3* |  | CCAACTACCGCGCTTATGC | CTCGCTCCAGCTCTTGCTTT |
| *SCN5A* |  | CACCAACTGCGTGTTCATGG | CAGAAGCCTCGAGCCAGAAT |
| *CPT1B* |  | CTCCTTTCCTTGCTGAGGTG | TCTCGCCTGCAATCATGTAG |
| *PDK4* |  | CCTTTGGCTGGTTTTGGTTA | CCTGCTTGGGATACACCAGT |
